# Lymph node swelling combined with temporary effector T cell retention aids T cell response in a model of adaptive immunity

**DOI:** 10.1101/2020.06.19.161232

**Authors:** Sarah C Johnson, Jennifer Frattolin, Lowell T. Edgar, Mohammad Jafarnejad, James E. Moore

## Abstract

Swelling of lymph nodes is commonly observed during the adaptive immune response, yet the impact on T cell trafficking and subsequent immune response is not well known. To better understand the effect of macro-scale alterations, we developed an agent-based model of the lymph node paracortex, describing T cell trafficking and response to antigen-presenting dendritic cells alongside swelling-induced changes in T cell recruitment and egress, and regulation of expression of egress-modulating T cell receptor Sphingosine-1-phosphate receptor-1. Analysis of effector T cell response under varying swelling conditions showed that swelling consistently aided T cell activation. However, subsequent effector CD8^+^ T cell production could be reduced if swelling occurred too early in the T cell proliferative phase or if T cell cognate frequency was low due to increased opportunity for T cell exit. Temporarily extending retention of newly differentiated effector T cells, mediated by Sphingosine-1-phosphate receptor-1 expression, mitigated any negative effects of swelling by allowing facilitation of activation to outweigh increased access to exit areas. These results suggest targeting temporary effector T cell retention and egress associated with swelling may offer new ways to modulate effector TC responses in, for example, immuno-suppressed patients and optimisation of vaccine design.

## Introduction

The lymphatic system is a network of organs and lymphatic vessels (LVs) that maintains fluid balance and delivers crucial antigen information to lymph nodes (LNs) for adaptive immunity initiation. LNs contain compartments populated by T cells (TCs), B cells, fibroreticular cells (FRCs), and lymphatic endothelial cells (LECs) [1, 2]. When antigens are presented (either suspended in lymph or captured by incoming antigen-presenting cells such as Dendritic Cells (DCs)), the LNs’ physical environment changes. Swelling of LNs is a well-known consequence of antigen presentation, but the effects of swelling on processes crucial for adaptive immunity are not well understood.

TCs and B cells mainly enter LNs by transmigrating from blood vessels in the paracortex, while lymph-borne DCs migrate into the paracortex across the sub-capsular-sinus (SCS) floor [3, 4]. Typically, 1 in 10,000 naïve TCs express a complementary TC receptor to the antigen fragment presented by DCs within a MHCI (to CD8^+^ TCs) or MHCII (CD4^+^ TCs) molecule [5, 6]. With sufficient affinity and stimuli, TCs undergo activation, secrete inflammatory and activation-facilitating cytokines and differentiate into effector and memory TCs [7]. The mechanisms driving LN swelling include DC presence, B cell signalling and trapping of non-activated TCs [8–11]. Regardless of the trigger, within 2 days, TC exit rate drops (“LN shutdown”), blood flow to the LN increases, and inflammatory signalling results in a 3-5 fold increase in TC recruitment via high endothelial venules (HEVs) [12–15]. From 48-96 hours, LN mass increases 2-5-fold, accompanied by a similar increase in cellularity, and FRCs elongate to accommodate LN size increase [11, 16, 17]. Subsequent LEC and FRC proliferation allows maintenance of LN architecture during further expansion [10, 17, 18]. The LN blood vessels also grow, increasing blood vessel volume roughly proportional to overall LN volume, accompanied by further TC recruitment [9, 14, 19]. Between 2- and 5-days post-immunisation, antigen-presenting DC number in the LNs peaks, TC activation and proliferation is underway and TC egress increases 3-6 fold [10, 11, 20, 21]. Expansion of medullary and SCS areas aids increased TC egress [22]. Recruitment of TCs then declines, HEV, FRC and TC proliferation subsides, remaining effector TCs may undergo apoptosis and LNs return to baseline volume while memory cells recirculate [19]. Throughout these processes, TC egress is modulated by Sphingosine-1-Phosphate-1 receptor (S1P_1_r) expression and chemokine signalling axes. After entering the LN, TCs express S1P_1_r) at low levels but begin S1P_1_r re-expression after 2 hours [23, 24]. TCs exit LNs by probing and subsequently entering cortical sinuses in the paracortex or the medullary interface, aided by chemotaxis [25, 26]. During inflammation, TC S1P_1_r expression is reciprocally regulated by CD69, an early TC activation marker. This mechanism contributes to the initial decrease in TC egress, termed LN shutdown, and later to the specific retention of activated TCs [15, 27]. Differentiated effector TCs re-express S1P_1_r, facilitating egress [28].

The ability to investigate the importance of LN swelling in these processes is limited experimentally by a lack of means to modulate swelling without interfering directly with other aspects of adaptive immunity. We chose to develop an Agent Based Model (ABM) that could describe macroscale geometric changes, microscale TC and DC interactions and capture emergent behaviour by modelling the probabilistic behaviour of thousands of cells. Beyond the desire for a better understanding, we aim to provide a means for designing experiments that explore potential therapeutic means of modulating LN swelling.

Fixed-volume ABMs have provided insight into interactions relevant to vaccine design, for example, the effects of antigenic peptide separation on TC activation, influential aspects of TC-DC interaction, and memory TC production [29–33]. An ABM to investigate chemotactic influence included a form of paracortical expansion, where grid compartment number remained equal to TC number and exit portal number altered to maintain a mean TC residence time. This model suggested relative chemokine level is important but may underestimate changes in crowding and egress with swelling [34–36]. Simulations integrating a fixed-volume lattice-based model and a continuous model of chemokine diffusion showed early antigen removal and TC exit regulation affected the balanced system dynamics, indicating that macroscale swelling is likely to significantly affect micro-scale TC activity [37].

In summary, the careful trafficking and coordination of immune cell movements in the LNs, suggests LN swelling may significantly impact the adaptive response. We developed a computational ABM to investigate this hypothesis. The results suggest an important role for regulating early effector TC retention to maintain the benefits of LN swelling on overall effector TC response.

### ABM geometry

We aimed to replicate a murine LN by integrating experimentally obtained parameters. The paracortex was modelled as a sphere with initial radius *R*_0_=200*µ*m, derived from confocal images of murine LNs [2, 38]. Geometric symmetry was assumed so that one-half of total spherical geometry was modelled. The modelling domain was divided into cuboid grid compartments, with edge length 6*µ*m (Fig 1C). For each grid compartment, we tracked which region of the paracortex was represented, such as ‘exit’, ‘boundary’ or ‘outside’.

**Figure 1:**
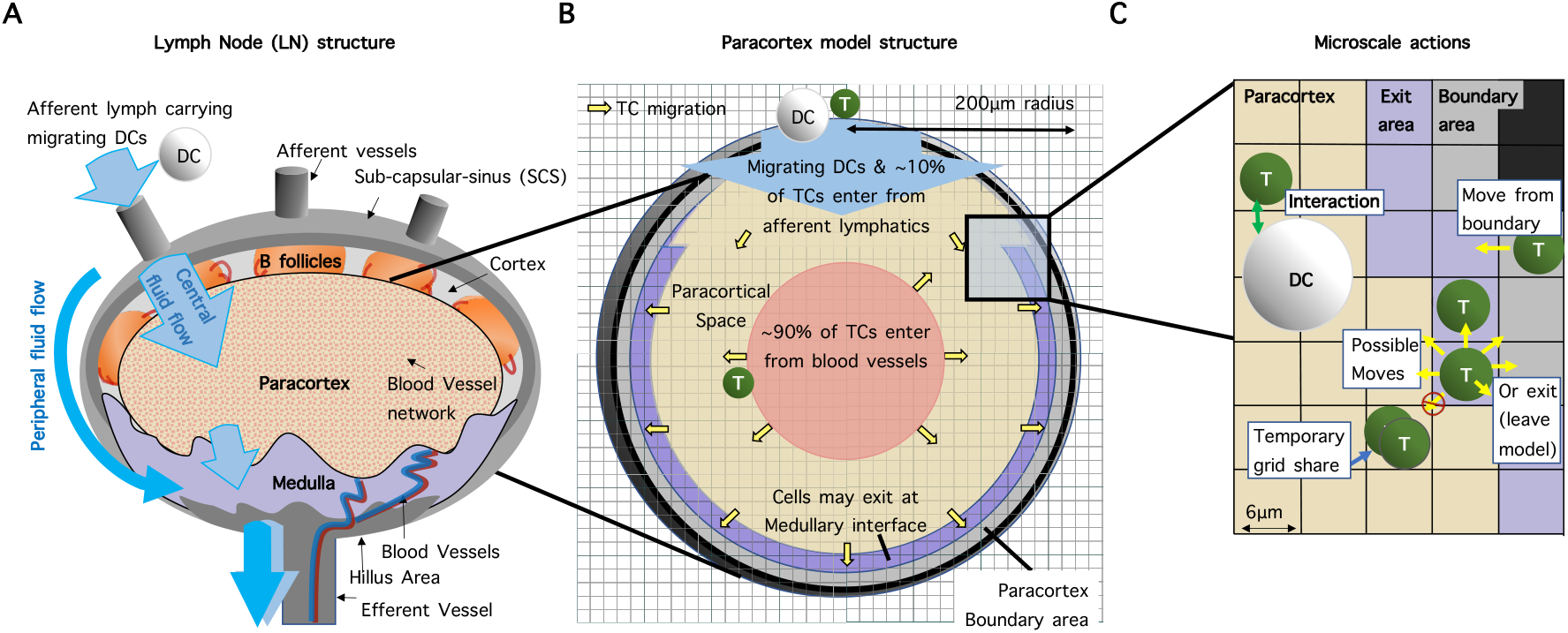
Model geometry and structure. (A) LN structure displaying arriving lymph containing agDCs. (B) TCs enter in the centre of the spherical paracortical model and exit near the interface with the medulla and SCS. The paracortex radius expands as a function of TCs present. (C) TCs move to adjacent grid-compartments, interact with neighbouring agents and are influenced by grid-compartment properties, which are updated each time-step.

### Modelling Swelling

Paracortical swelling or contraction was achieved by changing the region type that each grid compartment represented, while maintaining entry and exit areas as a percentage of the defined outer radius (Fig 2A). We collected data from murine experiments regarding change in LN mass and volume, TCs, structural cells, migrating DCs, TC recruitment and TC egress following antigenic stimulus application [9, 11, 12, 19, 20]. Based on these data we calculated paracortical volume as a sigmoidal function of TCs present (Fig A, S1 File). Initial TC increase is permitted without triggering significant swelling, reflecting initial inhibition of stromal cell proliferation by secretion of IFN type 1 [39]. A delayed volume increase in response to TC number is in agreement with the cell-signalling switch at day 2 to favour LN expansion, through mechanisms such as increased elasticity of the FRC network and LEC proliferation [11, 40].

**Figure 2:**
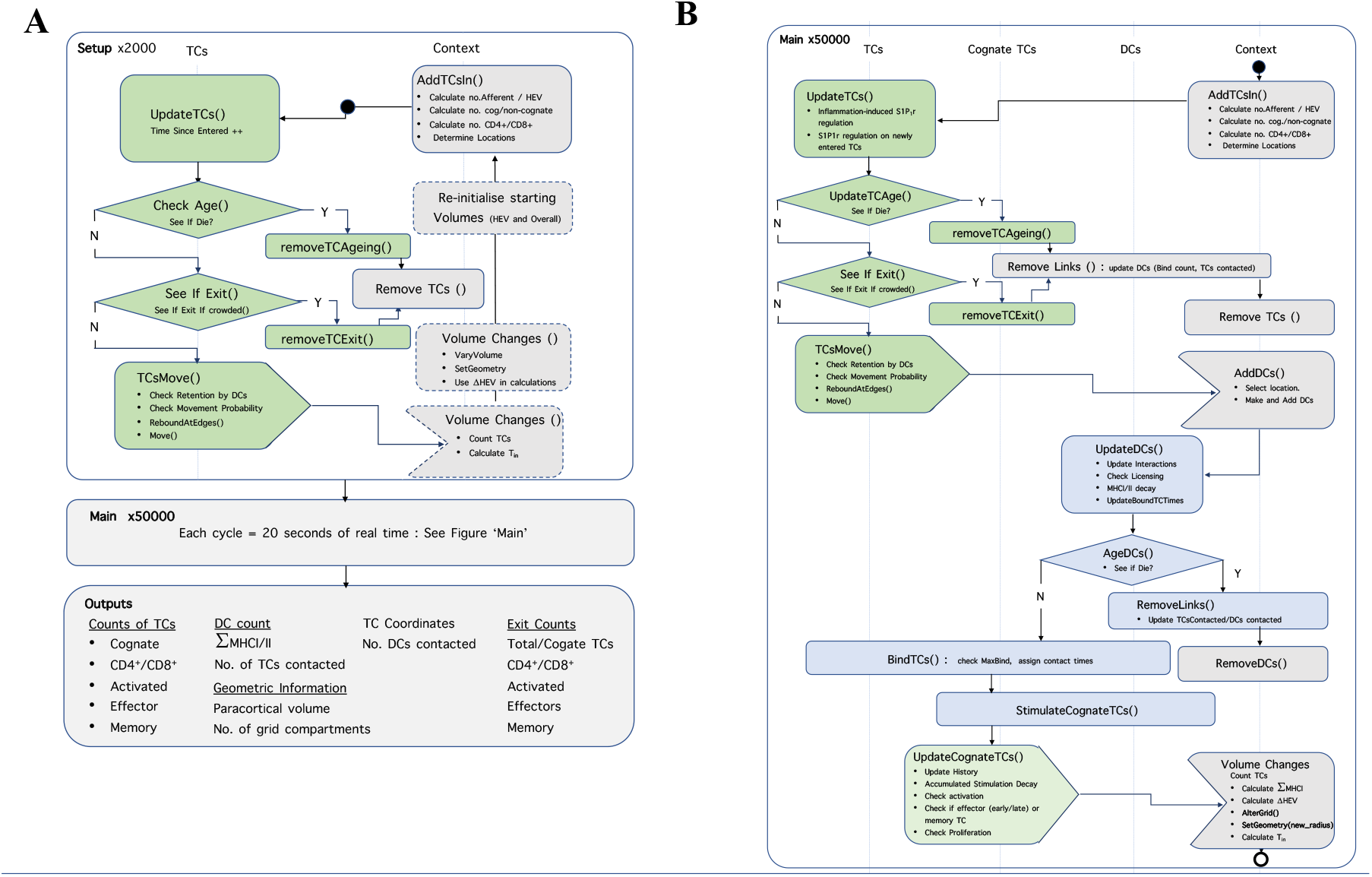
Structure of the model code. A) The model is initiated in the absence of stimulus, capacity for paracortical volume change is then introduced and variables storing starting volumes are updated. Agents represent cells (TCs and DCs), store interaction history and present state information. The ‘Context’ describes the environment and ‘Projections’ between agents allow information transfer. Each time-step represents 20 seconds. B) Following equilibration, the ‘main’ function calls a repeated series of sub-functions (see S2 File) describing DC arrival and TC response.

### TC recruitment

Under baseline conditions, TC recruitment rate was specified as 2000 TCs/hour, with naive TC transit time (T_*res*_) defined between 6-24 hours and a constant TC-to-compartment ratio assumed (1.2 in S1 File). In accordance with HEV images, 90% of TCs entered at ‘entry’ compartments designated as the inner half of the paracortical radius [41]. Remaining TCs entered via the SCS interface.

The volume of the entry grids was representative of blood vessel volume (*V*_*B*_), which changes proportionally with paracortical volume [9, 42]. We assumed TC recruitment rate (T_*in*_), is influenced by 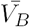 (Eq. 1). Acute recruitment changes due to inflammation-induced signalling cascades at the HEVs were represented by inflammatory index, I_*F*_ . This index affects TC influx when antigenic presence D (sum of MHCII, Eq. 2) rises above threshold, T1, the minimal DC number required to elicit a response [30]. The value of I_*F*_ increases proportionally with antigenic presence by a recruitment factor (R_*F*_) up to a maximum inflammation-induced TC recruitment, threshold T2 (Eq. 3). TC influx was therefore defined as:

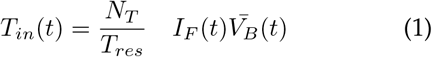

Where N_*T*_ is initial TC number, T_*res*_is naïve TC transit time, I_*F*_ is the inflammatory index, and 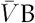 is normalised blood vessel volume. Threshold values for T1, T2 and R_*F*_ were estimated from initial TC recruitment rate changes due to inflammation, whilst considering changes due to HEV growth and agDC number present [11–15, 20, 43]. Inflammatory index I_*F*_ was calculated as:

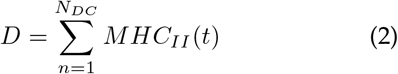

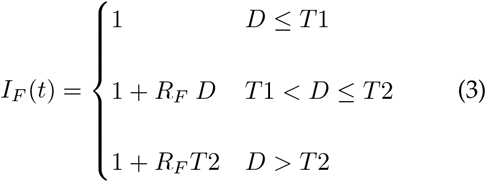

Where N_*DC*_ is number of agDCs present, D is the sum of MHCII carried by each agDC and Recruitment Factor R_*F*_ is an estimated increase in recruitment rate.

### TC egress and S1P_1_r expression

We altered relative TC expression of S1P_1_r (SP), from a default value of 1, under three conditions (Fig B in S1 File). Following TC entry into the paracortex S1P_1_r remained down-regulated (SP_*in*_=0.1) for 45-180 minutes, before re-expressing due to low paracortical S1P concentration [24]. A ‘LN shutdown’ mechanism was included by down-regulating S1P_1_r (SP_*inflam*_=0.4) on all TCs when sufficient antigenic presence (summation of MHCII) was detected, estimated to correspond to 6 hours post-agDC appearance. Activation-induced TC S1P_1_r down-regulation was represented by decreasing S1P_1_r expression 10-fold when TCs initially activated (SP_*act*_=0.01), increasing S1P_1_r expression as TCs differentiated into early effector TCs (SP_*early*_=0.4), and further increasing expression when effector TCs underwent *≥* 8 divisions (SP_*late*_=1) [23, 44, 45].

### TC and DC motility and interaction

DCs were modelled as 6*µ*m radius spheres and interacted with TCs within a two-grid radius, up to a maximum number of TCs at once (B_*max*_). TCs were modelled as spheres of volume 150*µ*m^3^, that initially occupied 60% of the total paracortex volume [46]. The frequency of antigen-specific (cognate) TCs (F_*cog*_) was derived from *in-vivo* reports [6]. Each agDC presented a decaying MHC signal, and during interactions cognate TCs gained ‘stimulation’ (S) at rate *κ*s, proportional to MHC presented, while losing stimulation at rate *λ*S. Similar to previous models, probability of TC activation and, after a minimum of 4 proliferation’s, differentiation into effector or memory TCs, was determined as a sigmoidal function of accumulated stimulation [31, 35, 47]. See S1 File for full rules.

### Computation

We build a class-based ABM (Fig C in S1 File) in java using RepastSimphony (repast.sourceforge.net) with repeated rules each time-step (Fig 2). Further UML descriptions are in S2 File. We carried out batch simulations on the Imperial College High Performance Computing cluster and analysed data in Matlab. Model code is available on GitHub at johnsara04/paracortex_model_johnson19.

### Parameter selection and sensitivity analysis

We estimated our parameters from published studies with inflammation-induced mice or previous relevant models (Table A in S1 File). To ensure awareness of influential but uncertain or biologically unconstrained parameters, we carried out a global sensitivity analysis. We used Latin Hypercube sampling to select 300 parameter combinations, simulated each set 3 times and recorded TC number (activated, effector, memory, effector exited and memory exited). Partial rank correlation coefficients (PRCCs) were calculated between each parameter and output for each day (3-13), assuming monotonic relationships [48]. We report significant PRCCs with strength greater than 0.2 (S4 File).

### Validation and model robustness

To ensure we did not overfit the model to one swelling scenario, we simulated four experiments that mimic *in-vivo* and/or *in-vitro* experiments, holding our parameter selection constant, aside from a single parameter. In each scenario, we compared the effects on TC activation and CD4^+^ and CD8^+^ effector TC response to relevant published studies. We inhibited S1P^1^r down-regulation on activated TCs as carried out by Gräler et al, and Lo et al [24, 49]. We varied the initial proportion of cognate TCs, as carried out by Obar et al and Moon et al [50, 51]. We varied agDCs number as carried out by Kaech et al, and Martín-Fontecha et al, and we simulated early DC apoptosis, as carried out by Prlic et al 2006. [52–54].

## Results

### The model produces realistic baseline TC motility and response to agDCs

We confirmed that the calibrated model produced an average TC velocity (n=200) of 13.1*µ*/min, reaching up to 24*µ*m/min (Fig 3A), in-line with murine *in-vivo* measurements [41, 55–58]. The mean TC paracortex transit time was 13.1 hours (n=16,000), ranging from 20 minutes to *>*60 hours (Fig 3B), in-line with observations that 74% of CD4^+^ TCs and 64% of CD8^+^ TCs transit murine LNs within a day [59]. The linear relationship between TC displacement versus square root of time, (Fig 3C) illustrated maintenance of random walk behaviour [60]. The motility coefficient (CM) was 63.2*µ*m^2^/min, which is within the 50-100*µ*m^2^/min range observed in mice [61].

**Figure 3:**
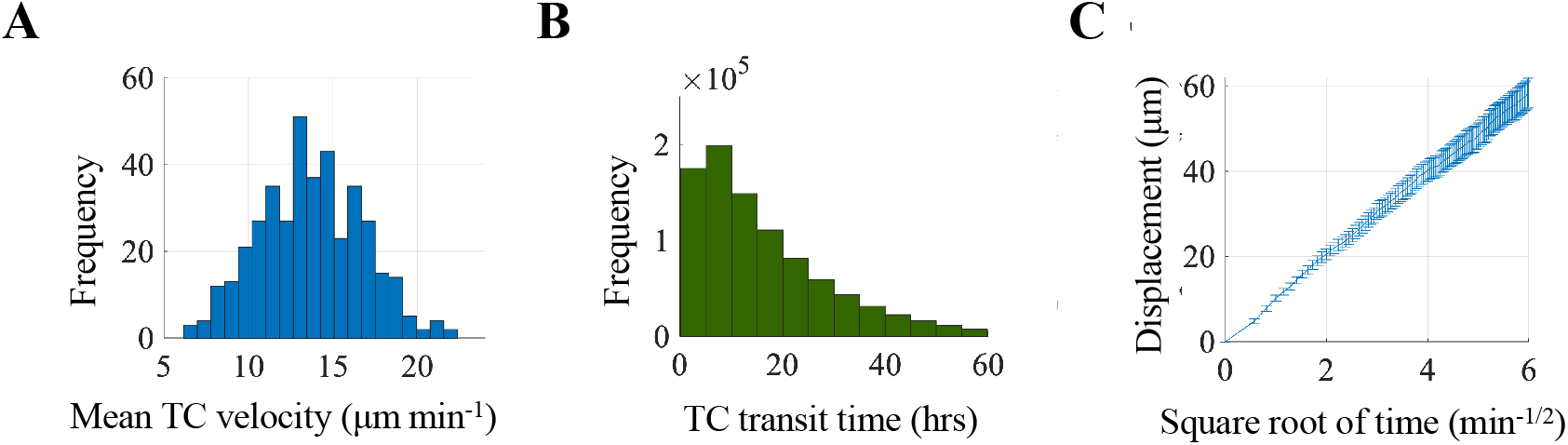
Baseline TC motility (n=200). A) Mean TC velocity. B) Most TCs transit in *<*24hrs. C) Mean (^+^/-SEM) of TC displacement showed a linear relationship to the square root of time, indicating random walk behaviour.

TC responses to agDC stimuli corresponded well to data from *in-vivo* trials in mice, sheep and rats, displaying the expected phases of TC trafficking and response (Fig A in S3 File). TC numbers began to increase approximately 6 hours after initial agDC entry, and by day 11 had returned to within 15% of pre-stimulus values (Fig 4B), in-line with temporal responses observed *in-vivo* [11,12,20]. The appearance of activated, effector, and memory TCs began at 16-24 hours, day 3.5 and day 5 post-agDC entry, respectively, in agreement with *in-vivo* reports and cell-culture models [62, 63]. Effector CD4^+^ TCs appeared 1-1.5 hours before CD8^+^ effector TCs (Fig 4H, I). As observed *in-vivo*, the peak cognate CD8^+^ TC number was an order of magnitude higher than that of CD4^+^ TCs [64, 65]. The contraction phase began at day 7 and continued through day 11 (Fig 4B). Increase in TC egress rate peaked a day later than the increase in TC entry rate (Fig 4F,J), corresponding well with *invivo* observations [16, 66].

**Figure 4:**
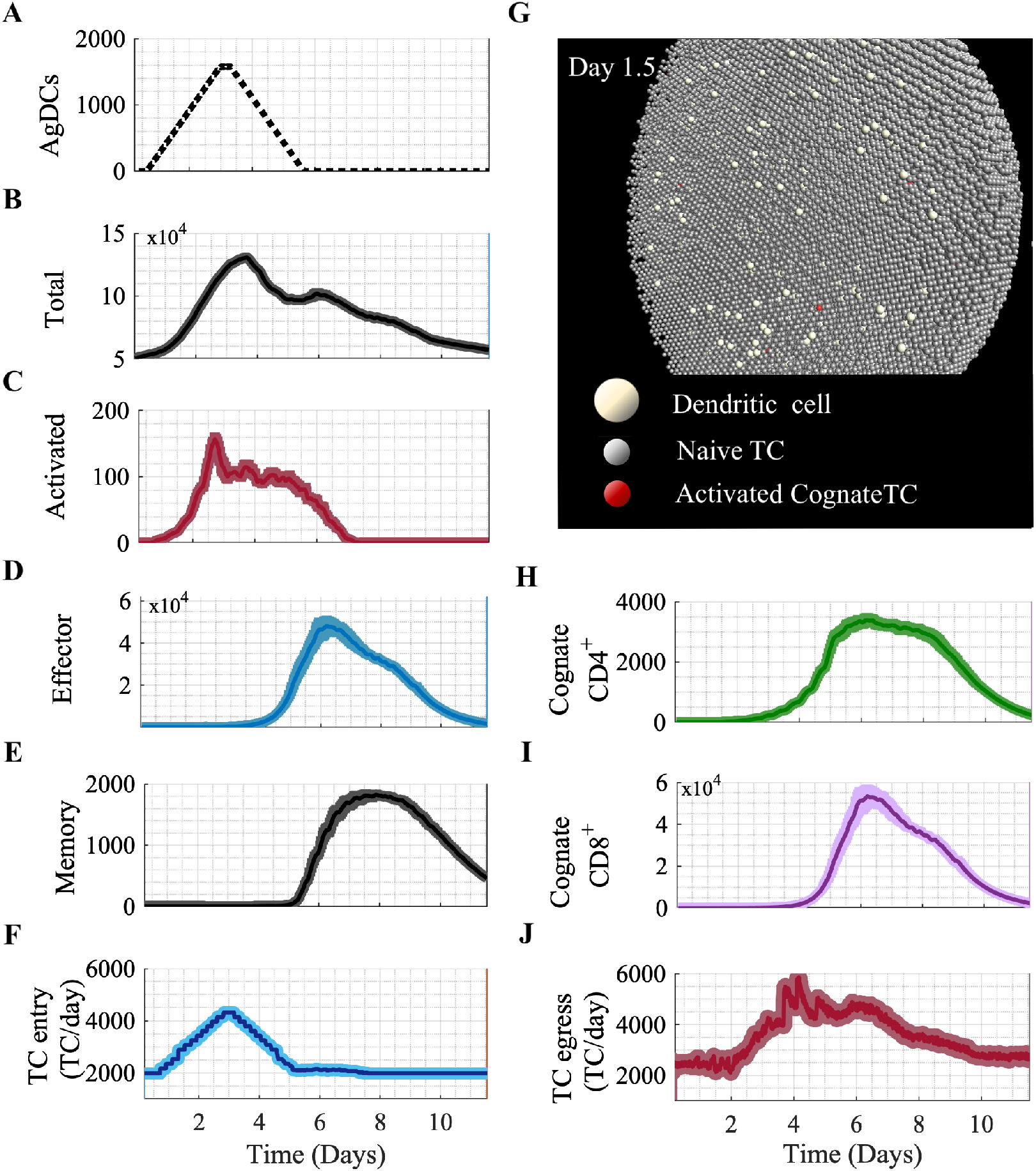
TC responses in the paracortex following entry of agDCs under baseline conditions. Average result with SEM of 12 simulations. A) Incoming agDCs. B) The total TCs number peaked at 3.5 days, comprising mainly of non-cognate naïve TCs. C) Activated TC appearance began 12 hours after the first agDCs entered. D) Effector TC number peaked at day 6. E. Memory TCs appeared at 5 days and 25% of the peak number remained at the simulation end. F) TC entry rate increased 2x, peaking at day 3. G. Model interface showing day 1.5 with agDCs present and TC activation initiated. H. Cognate CD4^+^ TCs began extensive proliferate at day 2.2. I. Cognate CD8^+^ TCs began proliferation at day 4 and reached numbers 10x more than cognate CD4^+^ TCs. J. TC egress rate declined between day 1 and 2, then increased 3x by day 4.

### Model robustness

Holding the default parameters and varying a single parameter at a time to mimic *in-vivo* and *in-vitro* experiments resulted in reasonable TC behaviour. For example, preventing S1P_1_r down-regulation post-antigenic stimulus detection *in-silico* reduced activated TC number by 60%-81% (Fig 5A). A study transferring activated TCs that over-express S1P_1_r into mice LNs, removing S1P_1_r-mediated retention, resulted in 90% less activated TC retention compared to control mice when measured 15 hours later (Fig 5B [24]. A study using mice with constitutive TC expression of S1P_1_r showed a 40% reduction in activated TCs post-immunisation (Fig 5C) [49]. See S3 File for comparison of varying cognate frequency, agDC presence and duration of stimuli application.

**Figure 5:**
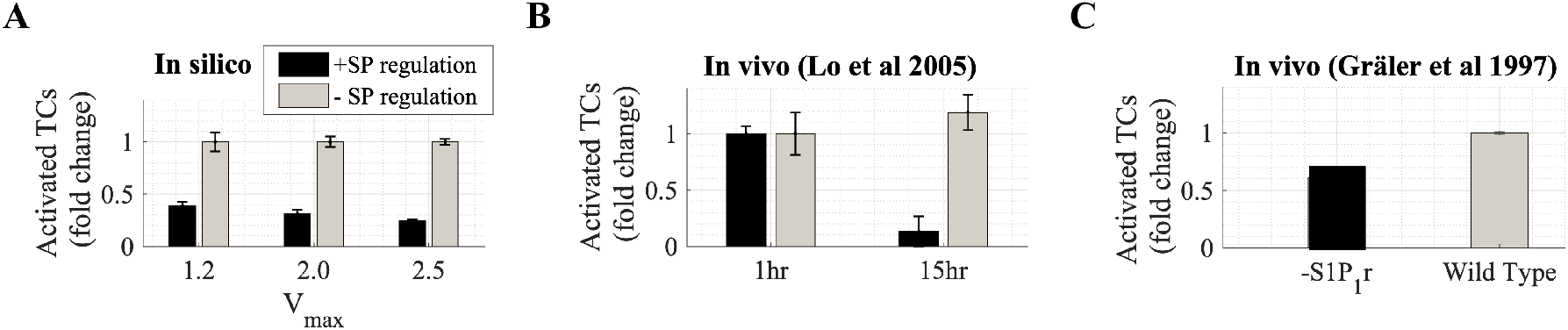
Comparing model predictions with reported *in-vivo* effects of S1P_1_r down-regulation on TC activation. A. Simulation results (n=10) with and without S1P_1_r downregulation (+/-SP regulation) showed total activated TCs number reduced 60%, 72% and 81% (Mean+/-SEM) at V_*max*_=1.2,1.5 & 2. B. Pre-activated TCs over-expressing S1P_1_r (-SP regulation) were transferred into mice. Retention of activated TCs 15 hours later fell by 90% compared to transferred wildtype TCs (SP+ regulation). Adapted from Lo et al 2005. C) In mice with constitutive S1P_1_r expression, activated TC number in LNs 24 hrs post-immunisation dropped by 40%. Adapted from Gräler et al 1997.

The global parameter sensitivity analysis indicated that the dominant parameters in determining the target outcomes of TC activation, total TC effectors, and TCs exited were F_*cog*_, T_*DCin*_, V_*max*_. The unconstrained parameters used to describe signal integration and parameterize activation or differentiation probability curves, were not identified as significantly influential in determining target outcomes (p*>*0.05, R^2^<0.2)(Fig A and Tables A-C in S4 File).

### Paracortical swelling consistently aids TC activation

When maximal swelling (V_*max*_) was varied from 1 to 2.8, activated TC number doubled and positively correlated with V_*max*_ (R^2^=0.96, p*<*10^*−*5^) (Fig 6A). The total number of effector TCs decreased by 15% (Fig 6B) and negatively correlated with V_*max*_ (R^2^=0.86, p*<*10^*−*3^) but the number of effector TCs that exited by day 10 did not significantly vary (Fig E in S3 File).

**Figure 6:**
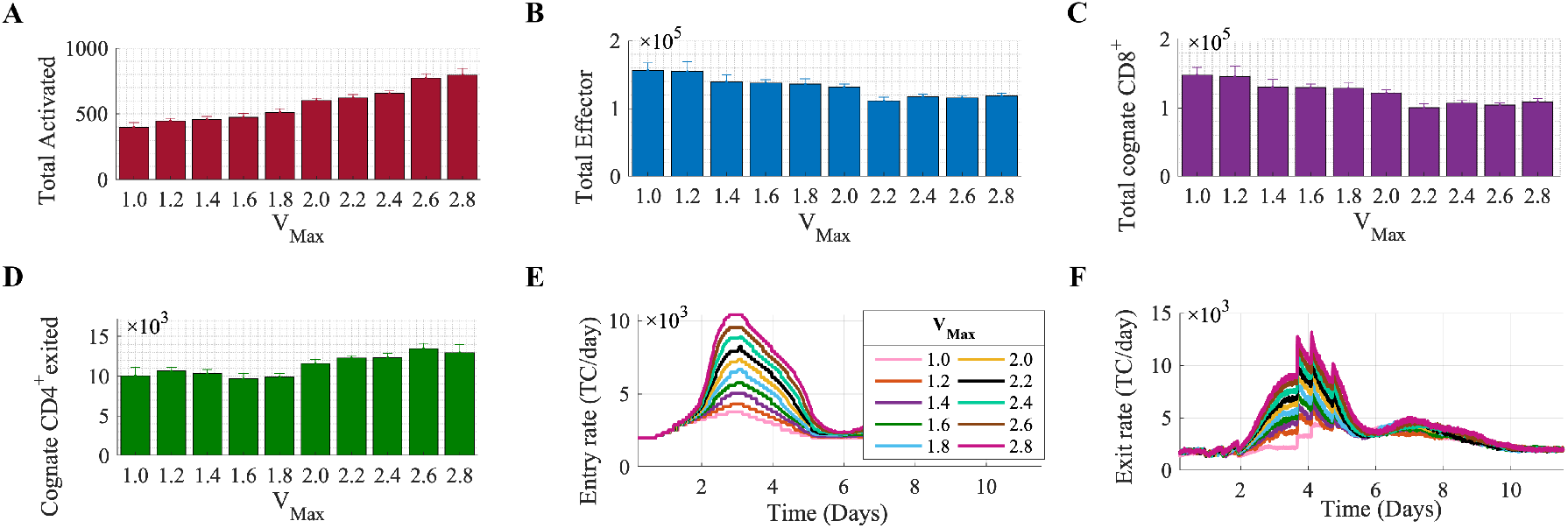
Changes in TC response in the paracortex when varying swelling. Between V_*max*_=1 and 2.8, A) total activated TCs doubled and positively correlated with V_*max*_ (R^2^= 0.96, p=1.07×10^*−*6^), B) total effector TCs decreased 0.3x, negatively correlating with V_*max*_ (R^2^=0.86, p=1.23×10^*−*4^), C) total cognate CD8^+^ TCs negatively correlated with V_*max*_ (R^2^=0.855, p=1.28×10^*−*4^), D) and total cognate CD4^+^ TCs that exited increased 1.3x, positively correlating with V_*max*_ (R^2^=0.76, p=0.001). E, F) Peak entry and exit rate increased proportionally to V_*max*_. Results are the mean of n *≥* 7 simulations with SEM displayed.

Assessment of TC subgroups showed that the total cognate CD8^+^ TCs present decreased by 25% (Fig 6C), negatively correlating with V_*max*_ (R^2^=0.855, p*<*10-3) but there was no change in the number of exiting cognate CD8^+^ TCs (Fig E in S3 File). Conversely, the number of cognate CD4^+^ TCs that left the paracortex by day 10 increased by 30% and positively correlated with V_*max*_ (R^2^=0.76, p=0.001) (Fig 6D) but cognate CD4^+^ TC present did not varied significantly (Fig E in S3 File).

The peak TC recruitment rate positively correlated with V_*max*_, meaning the absolute number of cognate TCs entering increased with swelling (Fig 6E). TC egress rate increased with V_*max*_ from day 3-6 (Fig 6F). Increased TC activation but decreased effector TC number remained when LN volume increased as a linear function of TCs (Fig A in S5 File).

### Reduced effector TC response with swelling was not due to a lack of agDC availability

We then analysed the mean number of interactions with DCs by cognate and non-cognate TCs present each day from day 1 to 6 at different maximal swelling (V_*max*_=1.20, 2.0 and 2.5). We found there was no decrease in the mean number of agDCs that each cognate TC contacted on all days (Fig 7A). We also found a slight increase in the number of contacts by day 3, a time-point that corresponds with peak swelling. The mean number of short contacts by non-cognate TCs decreases with swelling (Fig 7B). These results suggest there is no decrease in availability of DCs to cognate cells with swelling.

**Figure 7:**
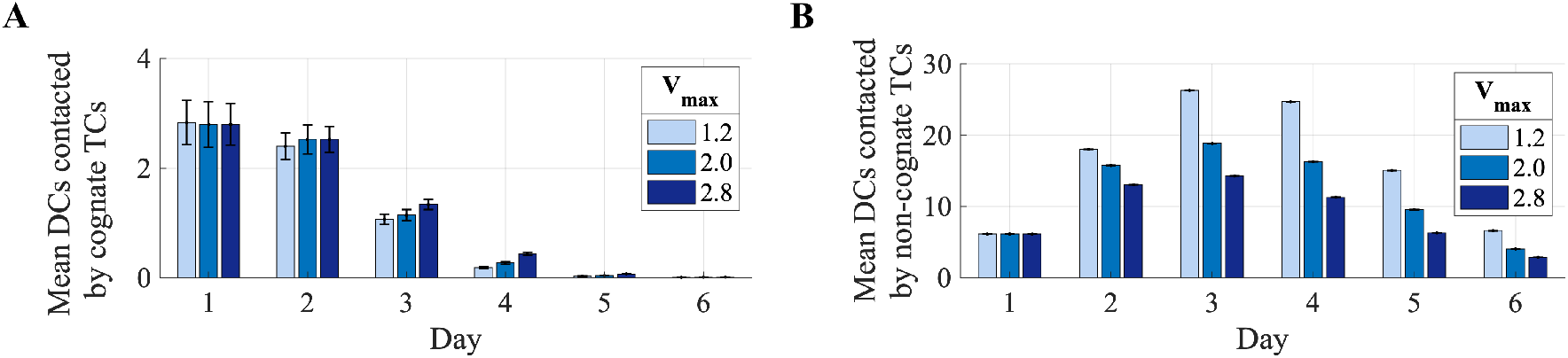
Changes in DC and TC contact with swelling from day 1 to 6. With increased swelling the mean number of DCs contacted by (A) cognate TCs did not decrease but (B) non-cognate TCs contacted less DCs.

### Paracortical swelling can hinder effector TC production in some circumstances

We carried out simulations with a small or large maximal swelling (V_*max*_=1.2 or 2.5) while applying a lower (8×10^4^ TCs) or higher (13×10^4^ TCs) T_*mid*_, making swelling occur relatively earlier or later (Fig 8A). Regardless of T_*mid*_ value, at least 40% more activated TCs were recorded with a large V_*max*_ compared to a small V_*max*_ (Fig 8B). With an earlier (low T_*mid*_) and larger swelling, the total number of effector TCs and effector TCs exited dropped significantly (p<0.05) (Fig 8C). However, with later swelling, (high T_*mid*_), a larger swelling no longer reduced effector TC number. This altered effector TC response was due to change in cognate CD8^+^ TC number, which showed the same pattern of results (Fig 8E). There was no change associated with T_*mid*_ in cognate CD4^+^ TC response (Fig 8F). Varying maximal swelling and T_*mid*_ over a wider range showed that the positive correlation of T_*mid*_ with effector TCs exited was only significant with a larger swelling (V_*max*_=2.5) (Fig 8D), likely due to the greater impact of varying T_*mid*_ with larger swelling (Fig 8A).

**Figure 8:**
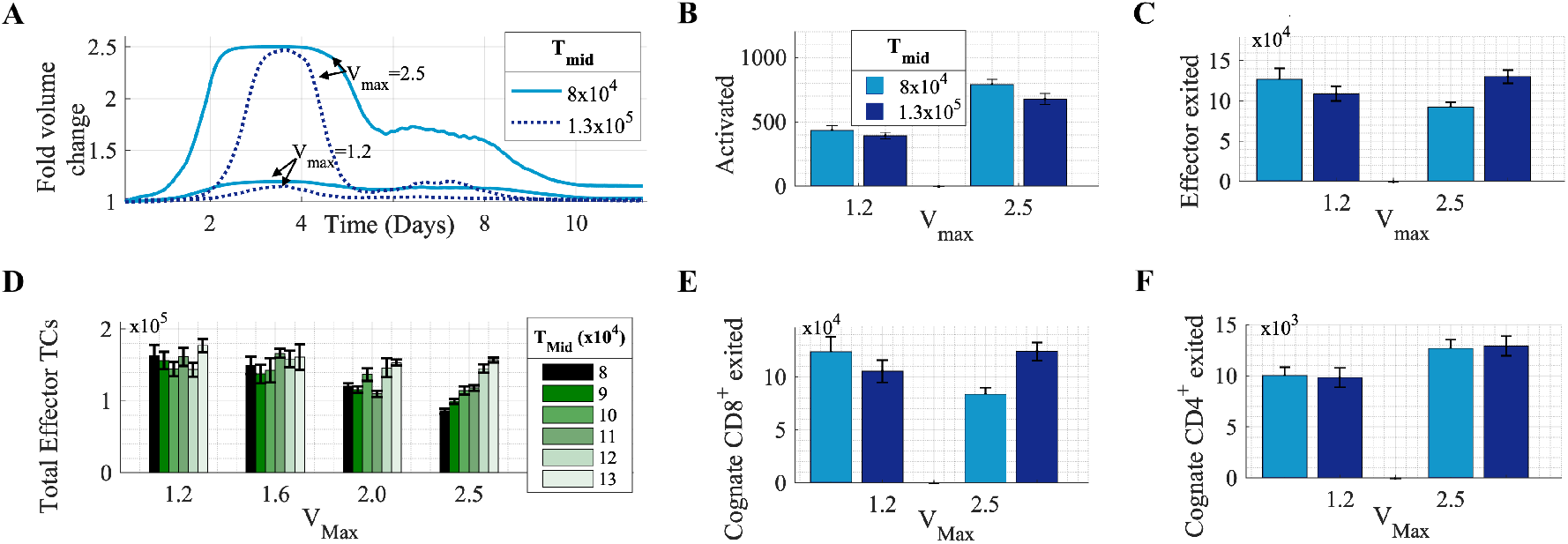
Varying the pattern of paracortical swelling. (A) The paracortex swells earlier and for a longer duration with a low T_*mid*_. (B) Increased swelling aided TC activation. C) With a lower T_*mid*_, as swelling increased, number of effector TCs exited decreased. (D) Further simulations varying T_*mid*_ confirmed that at a large swelling (V_*max*_=2.5), delayed swelling with a higher T_*mid*_ resulted in more total effectors TCs (R^2^=0.97,p= 3×10^*−*4^). (E) This effect was due to altered CD8^+^ TC number as (F) CD4^+^ TC number increased with V_*max*_ but was unaffected by varying T_*mid*_.

### S1P_1_r-mediated temporary retention of early effector TCs increased TC response

When we increased S1P_1_r down-regulation by lowering SP_*early*_ from the estimated default value of 0.4, a sustained increase in total TCs resulted, despite the action acting on early effector TCs only (Fig 9A). Unlike during simulations with default SP_*early*_ (Fig 6), effector TC number did not decrease with swelling. Instead, when SP_*early*_ was lowered from 0.4 to 0.1, approximately 15% and 10% more effector TCs were produced with larger V_*max*_ of 2.0 and 2.5 respectively. At every maximal swelling value SP_*early*_ inversely correlated with effector TC number (R^2^=0.92,0.93,0.92, p*<*0.005). Reducing SP_*early*_ from 0.4 to 0.05 doubled the number of effectors TCs exiting and increasing SP_*early*_ to 0.8 halved the number (Fig 9B).

**Figure 9:**
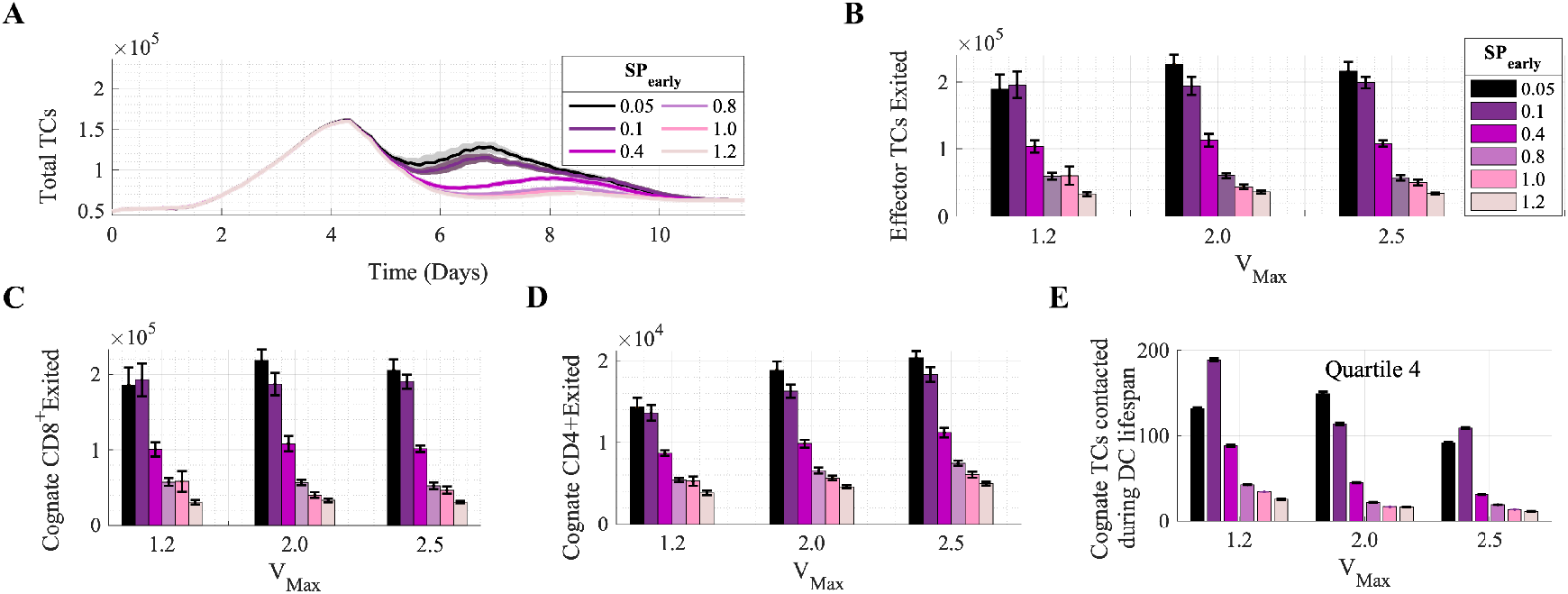
Temporary retention of effector TCs by modulating S1P_1_r expression on newly differentiated TCs (SP_*early*_). A) Reducing SP_*early*_ resulted in a higher total TC number. B) Number of effector TCs exited was affected more by SP_*early*_ than by V_*max*_, negatively correlating with SP_*early*_ (R^2^ *>*=0.92, p*<*0.005). SP_*early*_ negatively correlated with (C) CD8^+^ TCs exited (R^2^ *>*0.92, p*<*0.005) and (D) CD4^+^ TCs exited (R^2^ *>*0.91, p*<*0.005).E). The mean number of cognate TCs contacted increased as SP_*early*_ was lowered to 0.1 at each V_*max*_ but overall decreased with V_*max*_.

When analysing the TC sub-populations, both CD4^+^ and CD8^+^ effector TCs that exited the paracortex by day 10 doubled when SP_*early*_ was decreased from 0.4 to 0.05 (Fig 9C, D). This indicated CD4_+_ TCs do maintain further proliferative capacity in the model.

The number of TCs contacted by DCs increased as SP_*early*_ was decreased but overall decreased with swelling, therefore was not a driving factor of increased effector TC number (Fig 9E) implementation of an alternative model with non-specific constraint of TC egress by reducing expansion in the exit area also resulted in increased effector TC exit but produced unrealistic prolonged swelling above a 1.4-fold swelling (Fig B in S5 File)

### Non-specific early LN shutdown with a doubling of LN volume did not significantly impact effector TC production

We also varied the degree of initial LN shutdown, by varying SP_*inflam*_ from 0.1 (90% down-regulation) to SP_*inflam*_=1 (no shutdown). We permitted a doubling of LN volume. Increasing non-specific S1P_1_r down-regulation from 60 to 90% resulted in a sharp, 3-fold higher peak in the total number of TCs (Fig 10A), which is less physiologically realistic than with our default parameters. As SP_*inflam*_ decreased, TC activation increased (R^2^=0.83, p=0.01) (Fig 10B), but no trend with total effector TCs was identified (Fig 10C). We found no correlation between increased LN shutdown and the mean number of contacts with DCs by cognate TCs present at day 3 but a positive correlation with DCs contacted by non-cognate TCs (R^2^=0.93, p=0.0017) (Fig 10D).

**Figure 10:**
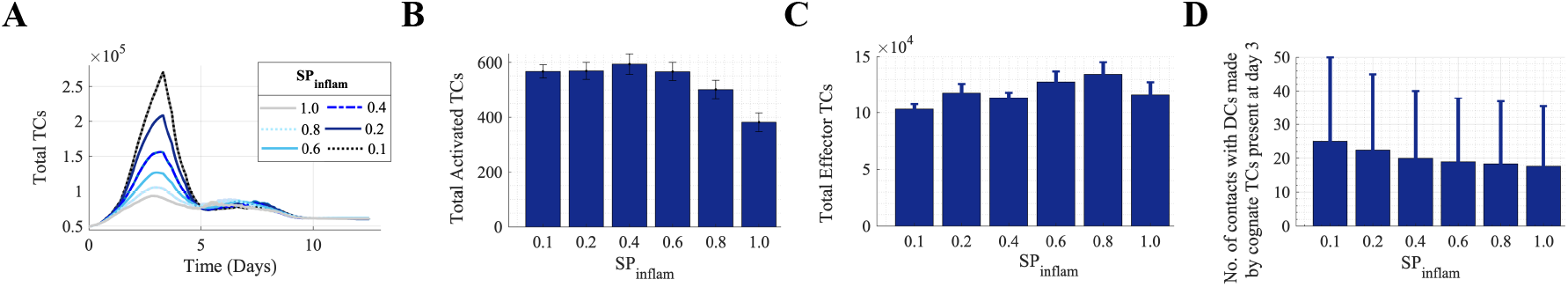
Varying LN shutdown by modulating initial inflammation-induced S1P_1_r downregulation (SP_*inflam*_). Modulating S1P_1_r expression from 0 to 90% downregulation (SP_*inflam*_=1 to SP_*inflam*_=0.1). A) total TC number decreases several fold but (B) TC activation increases (R^2^= 0.83, p=0.01) while (C) effector TC production shows no trend. D) Non-cognate TCs contact more DCs as SP_*inflam*_ is increased (R^2^=0.93, p=1.7×10^*−*3^).

### Boosting TC response when cognate TC frequency is low

Simulations using a 10-fold lower cognate TC frequency showed a larger decrease in effector TC number with swelling compared to the simulations with default cognition. With lower cognition, we observed a mean 73% fall with a 2-fold swelling, compared to a mean 17% decrease with 10-fold higher cognition (Fig 6B).

With V_*max*_=2.5, we recorded a mean 33% fall compared to a 5% decrease with 10-fold higher cognition (Fig E-vii in S3 File). We repeated the simulations with increased early effector TC S1P_1_r down-regulation (SP(_*early*_=0.1). This resulted in swelling of 2.0 or 2.5-fold benefiting the response. Assessment of TC and DC interactions showed that this was not due to increase in contact with DCs (Fig 11C).

**Figure 11:**
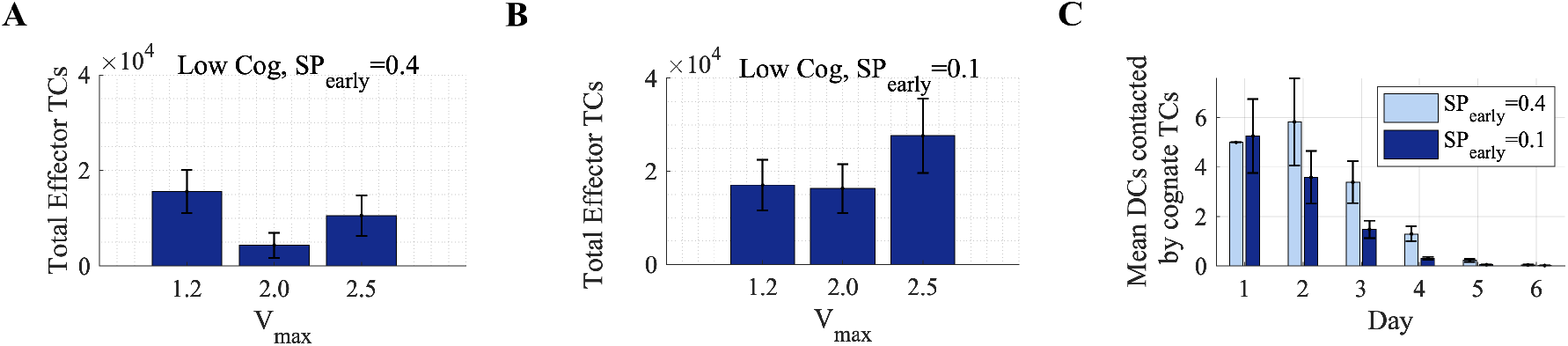
Swelling combined with increased retention of early effector TCs can improve response. With a 10-fold lower cognate frequency of 10^5^ and (A) default estimated S1P_1_r expression compared to (B) increased early effector S1P_1_r down-regulation. C. The increased response is not due to increased DC access (V_*max*_=2.0).

## Discussion

In this work, we aimed to better understand the effects of lymph node swelling in the formation of TC responses and identify key features that can influence TC behaviour. Our study builds on work using ABMs to investigate the impact of signal integration kinetics, TC migration and interaction dynamics on TC response with a focus on macroscale alterations and accompanying changes in egress and recruitment [29–31,33, 35].

We found that LN swelling consistently aids TC activation but increased swelling can inhibit subsequent effector TC response if it increased opportunity for effector TCs to egress prior to optimal proliferation. LN swelling aided increased TC recruitment and therefore a higher absolute number of cognate TCs in the paracortex, increasing TC activation probability in agreement with *in-vivo* TC recruitment studies [14]. In our model, swelling also presented a greater number of exit points and therefore increased opportunity for effector TC egress. Change in contact between TCs and cognate TCs was not a driving factor.

A key finding was that temporary S1P_1_r-mediated retention of newly differentiated effector TCs could increase effector TC production in scenarios where effectors egress prior to reaching sufficient proliferation. The increased production was not due to increased contact with DCs (Fig 9E), and non-specific TC retention in the first few days had no impact on effector TC response (Fig 10). Swelling could also increase effector TC production when the exit area growth with swelling was constrained in alternative models (Fig B in S5 File).

We also found that, with low TC cognition rate, temporary S1P_1_r-mediated retention of newly differentiated effector TCs could double effector TC response when combined with swelling, but swelling alone negatively impacted response. Here, swelling can increase initial TC recruitment and initial absolute number of cognate TCs but must be combined with increased temporary retention of newly differentiated cells to benefit response.

The temporary nature of this S1P_1_r modulation is crucial to increase effector TC number. Permanent inhibition of effector TC S1P_1_r expression has been carried out *in-vivo*, and therapeutically, S1P_1_r down-regulation is the mechanism of multiple sclerosis drug Fingolipid. This acts to indefinitely retain effector TCs in the LN to prevent autoimmune response [67]. Temporary down-regulation on selectively newly-differentiated TCs may prove technically difficult, suggesting alternative means of retention is desirable [28].

In contrast to our results, transferring 10^6^ cognate TCs into murine LNs, whilst facilitating swelling by inducing FRC elongation and inhibiting FRC contraction enhanced subsequent TC proliferative response [17]. The authors suggest this may be due to reduced inhibition of TC activation by FRCs, or increased DC migration. Despite omission of these features, our model is in agreement with the increased TC activation. With an inflated initial cognate TC number that exponentially proliferated, the proportional effects of TC egress may be less as TC proliferative response is also relative to starting cognate TC frequency (Fig B in S3 File) [6, 50, 51]. We may also overestimate the negative effects of egress area availability with swelling, but highlighting the sensitivity of egress changes and temporary retention as a means to counteract sub-optimal responses remains an important result.

Our model contains unconstrained parameters that relate to signal gain and loss (*κ*s and *γ*), activation and differentiation probability curves (Act*µ*4^+^, Act*µ*8^+^, Dif*µ*4^+^, Dif*µ*8^+^), TC recruitment and paracortical swelling (S4 File). The sensitivity analysis showed that the parameters populating activation and probability curves were not highly influential, and the influence of signal integration patterns has been the focus of previous modelling studies [31, 35, 47]. Our model was not over-fit to a single scenario as we also varied stimuli strength, duration and TC cognition rate with results correlating well with comparable *in-vivo* experiments.

Limitations of our model include lack of chemotactic influences, FRC network omission, and a simplified LN geometry. We omitted DC migration and LN-resident DCs, but our results indicate DC availability is not a limiting factor. We prioritised inclusion of S1P_1_r down-regulation over the role of chemokine CCR7 because when CCR7 and S1P_1_r TC expression is inhibited *in-vivo*, TCs migrate to the paracortex boundary but lack of S1P_1_r expression prevents exit [28]. The critical influence of retention in our model suggests future iterations should include a wider range of retentive influences.

Several models suggest TC contacts are not significantly influenced by FRC network inclusion and we assumed that, regardless of the underlying FRC structure, TCs migrate with a random walk [55, 58, 61, 68–70]. When the FRC is modelled as a small-world network, damaging the network by removing 50% of nodes can significantly affect effector TC response [71]. We assumed that FRC stretch and proliferation helps maintain FRC architecture during our modest swelling [11].

Model fidelity is also limited by a lack of information on exit-point availability during swelling, but the sensitivity to alterations in egress suggests that exit area change with swelling presents as a crucial area to focus future studies. Future model iterations including features, such as lymph flow and pressure alterations (along with fluid exchange with nodal blood vessels), could also significantly improve the representation of swelling, and thus TC egress and retention. It has been well-established that changes in hydrostatic and oncotic pressure differences across nodal blood vessel walls can reverse the net fluid exchange [72, 73]. Afferent lymphatic flow, and thus DC number, to LNs also increases with immune response, as well as influencing chemokine concentration fields and likely mechanoresponsive cell expression of signalling molecules and receptors. A key next step is therefore to couple the ABM to a computational flow model.

## Conclusion

Our results suggest that, although LN swelling aids TC activation, events that increase opportunity for TC egress prior to optimal proliferation, such as early LN swelling, can inhibit effector response. We found that temporary retention of newly-differentiated effector TCs may boost effector TC response. This effect is particularly of interest when initial TC response is small, for example, in immuno-suppressed patients, or desirable, such as when optimizing vaccine design to minimise antigen dose. Although permanent blockade of effector TC egress has been utilised to treat multiple sclerosis, temporary retention of effector TCs to boost subsequent effector TC production presents as a novel mechanism. Our model identified the importance of alterations in TC egress with swelling and emphasises the influence that retentive features, including factors such as chemokines, may have on effector TC response, which may be more practical *in-vivo* targets to manipulate.

## Supporting information

**S1 File. Supplementary Methods A**. 1.1 LN swelling, 1.2 TC recruitment, 1.3 Agent migration, 1.4 Agent interaction and signal integration. Parameter Tables.

**S2 File. Supplementary Methods B**. UML diagrams.

**S3 File**.**Supplementary Results A**. Model calibration (Fig A), model validation (Fig B-D) and LN swelling results (Fig E).

**S4 File**.**Supplementary Results B. Global sensitivity analysis**. Fig A & Tables A-C. Outputs are activated, effector and memory TCs present and effector and memory TCs exited.

**S5 File. Supplementary Results C**. Alternative models. 5.1 & Fig A. A model with linear TC to LN volume relationship. 5.2 & Fig B. Alternative models of TC crowding and egress.

## Competing Interests

We declare we have no competing interests.

## Funding

This study was supported by the Royal Society, The Royal Academy of Engineering, The Sir Leon Bagrit Trust and the U.S. National Institutes of Health (NIH) Grant U01-HL-123420.

## Acknowledgements

The authors gratefully acknowledge the assistance provided by Dr Samira Jamalian and Willy Bonneuil.

## Notes

https://github.com/johnsara04/ABM_Supplementary_Figures

### Competing Interest Statement

The authors have declared no competing interest.

### Summary of Updates

Major amendments including how swelling may increase effector T cell egress probability and that temporary retention of effector T cells combined with lymph node swelling can aid overall effector T cell response. Supplementary data has been updated and reorganised.

