## Supplementary Files 1-5 for "Lymph node swelling combined with temporary effector T cell retention aids T cell response in a model of adaptive immunity": S1_File.pdf

### Supplementary File 1 : Supplementary Methods A

#### 1.1 LN swelling

**Table A** Data regarding changes in lymphocyte counts and proliferation, LN mass, LN volume, blood flow and stromal cell counts and proliferation were collected from a range of sources and analysed to identify common relationships between T cells and LN volume.

| Description | Stimuli | Model | Reference |
|---|---|---|---|
| LN mass, T cells, B cells, lymphocytes, total cellularity, Blood Endothelial Cells, migratory DCs, FRCs | Hindpad CFA/OVA injection | murine | [1] |
| LN volume, HEV length, LN images | Hindpad CFA/OVA injection | murine | [2] |
| LN volume, HEV length, B cell volume, | Hindpad LCMV injection | murine | [3] |
| LN images, Dendritic cells, endothelial cell proliferation, total LN cellularity, HEV proliferation | LPS-matured BMDC injection | murine | [4] |
| LN mass, peripheral blood T cells ($CD4^+$ /$CD8^+$), lymphocytes, T cell $CD4^+$, T cell $CD8^+$ | skin-application of dinitrofluorobenzene | murine | [5] |
| LN blood flow, LN weight, lymphocyte influx, HEV proliferation, cell proliferation | sheep erythrocytes | rat | [6] |
| LN total cell count, blood flow T cells (CD4+/CD8+), B cells, T cell proliferation, B cell proliferation | HCpG/LPS +/- OVA also HSV | murine | [7] |
| LN cellularity, FRCs, LECs, BECs, T cells (CD4+/CD8+), FRC/LEC/BEC proliferation | OVA/Mont-immunized | murine | [8] |

#### 1.2 TC recruitment

Under non-inflammatory conditions, it was assumed that TC entry and exit remain constant and TCs occupy a constant percentage (55%) of the total paracortical volume. The model represents a LN of 0.113-0.268 $mm^3$, implying a LN mass of 0.18-0.44mg (based on collaborative unpublished measurements of murine popliteal LN mass versus volume). Reported lymphocyte recruitment for a popliteal LN of 1.15g is $4x10^7$ lymphocytes/hour and up to 40% of lymphocytes are B cells [9–11]. The model represents half a paracortex, therefore TC recruitment rate was estimated as

1950-9000 TCs/hour under non-antigenic conditions. Naive TC transit time through the LN ($T_{res}$) was estimated as 6-24 hours [12].

#### 1.3 Agents and agent migration

The agents were designated as members of the TC or DC class (Fig A). The starting TC population is composed of 70% helper TCs ($CD4^+$) and 30% cytotoxic TCs ($CD8^+$) [13]. Each timestep ($\delta$t=20s), TCs can move one grid length to an available neighbouring grid compartment, moving with probability, $\beta$, and where availability is governed by crowding parameter $\gamma$. TC migration was assumed to follow a random walk with pauses. Previous models have described TC migration with Brownian motion, a random-walk with persistence,run and tumble, and Lévy walks amongst other methods, partly due to differing reports of *in-vivo* migration. [14–17].

Simulated DCs appear in the paracortex at a constant entry rate for 2 days, subsiding over the following 12 hours. Entry rate of DCs was scaled from counts of migrating DCs and initial TCs in a murine LN post-immunisation, with DCs totalling 4% of cells present [1].

#### 1.4 Agent interaction and signal integration

Interaction times with agDCs are drawn from uniform probability distributions with a mean of 3 minutes for non-cognate TCs ($T_{NC}$). Cognate TCs progress from 10-15 minute short interactions ($T_{short}$) to longer 50-70 minute interactions ($T_{long}$). During interactions TCs remain stationary.

Antigenic signal is presented by the DCs in the form of representative values of MHCI or MHCII (), and decays with time (t) with the form :

$$MHCI(t) = MHCI_i(0.5)^{\overline{\frac{t}{MHCI_{\frac{1}{2}}}}} \tag{1}$$

$MHCII_i$ is the initial MHCI or initial MHCII presented with respective MHC half-lives $MHCI_{1/2}$ and $MHCII_{1/2}$ estimated from *in-vitro* labelling of MHC molecules [18–21].

During cognate TC-DC interaction, $CD4^+$/$CD8^+$ TCs gain (S) at rate $\kappa$s while losing stimulation at rate $\lambda$S (Fig B.iii).Total change in TC stimulation is therefore given according to the first order rate equation:

$$\frac{dS}{dt} = K_s\, MHCII(t) - \lambda_S\, S(t) \tag{2}$$

Accumulated simulation decays to a minimal value of S=1 to allow differentiation between cognate TCs that gain and lose simulation (S=1) and those that never gain simulation (S=0). Probability of cognate $CD_{4+}$activation ($P_{a4+}$) or cognate $CD_{8+}$activation ($P_{a8+}$) is calculated with a sigmoidal function given by:

$$P_{a4^+} = \frac{1}{1 + e^{\frac{-S - Act\mu_4}{Actl_4}}} \tag{3}$$

Parameter $Act\mu_4$ determines the value of S required for 50% probability of activation and $Actl_4$ determines steepness of sigmoid inflection (Fig A.iv). Activation probability for $CD8^+$ TCs ($P_{a8^+}$) is determined with a sigmoid curve using a lower inflection point, ($Act\mu_8$), than for $CD4^+$ TCs. However, if the DC is 'licenced', which occurs post-interaction with an activated $CD4^+$ TC, $CD8^+$ TC and $CD4^+$ TC stimulation requirements are equal. This is to reflect facilitated $CD8^+$ activation as a result of activated CD4$^+$ induced production of cytokines [22]. Model parameters were estimated such that TC activation became apparent 8-15 hours post DC-arrival [23–25]. This method of signal integration and the subsequent progressive patterns of TC proliferation and differentiation is supported by *in-vivo* observations and modelling descriptions [25–28]. It is assumed that co-stimulatory requirements are met as agDCs are highly efficient antigen-presenting cells.

Post-activation, proliferation is possible every 11 (CD4$^+$) or 9 (CD8$^+$) ±1hr [29–32]. Differentiation into effector or memory cells is possible after >=4 divisions, with differentiation probability determined with a second set of sigmoidal probability curves with midpoint Dif$\mu_{4+}$ and Dif$\mu_{8+}$ respectively [33,34]. Greater CD4$^+$ TCs dependence on continued stimulation for differentiation than CD8$^+$ TCs was implemented by using a higher minimum threshold of accumulated stimulation for CD4$^+$ differentiation than for CD8$^+$ TCs [35–39]. The fraction of effector TCs that differentiate into memory TCs increases from 0.01 to 0.04 as TCs progress from 'early effectors' ($<8$ proliferations) to 'late effectors' [40]. This was implemented by assigning differentiation ratios of dif$_{early}$ and dif$_{late}$ to the two different subsets.

During TC activation and proliferation, S1P$_1$r expression was estimated from several *in-silico* studies. Expression change was considered following TC migration velocity into areas of high S1P concentration, changes in TC egress from the LN and changes in S1P$_1$R expression relative to naive S1P$_1$r expression following TC activation and with subsequent proliferation (Fig 2C.iii) [41–43].

**Fig A Modelling Methods.** (i). Definitions of different areas of the paracortex are based on the overall paracortical radius, and therefore alter in volume during expansion. (ii). During interactions, TCs gain stimulation at a rate proportional to the presented antigenic signal, which is itself decaying. Accumulated TC simulation also undergoes constant decay. (iii). Accumulated stimulation is to determine probability of TC activation or differentiation, dependent on the satisfaction of other criteria.(iv) The progression of TC proliferation and differentiation into effector and long-lived memory TCs with sufficient stimulation.

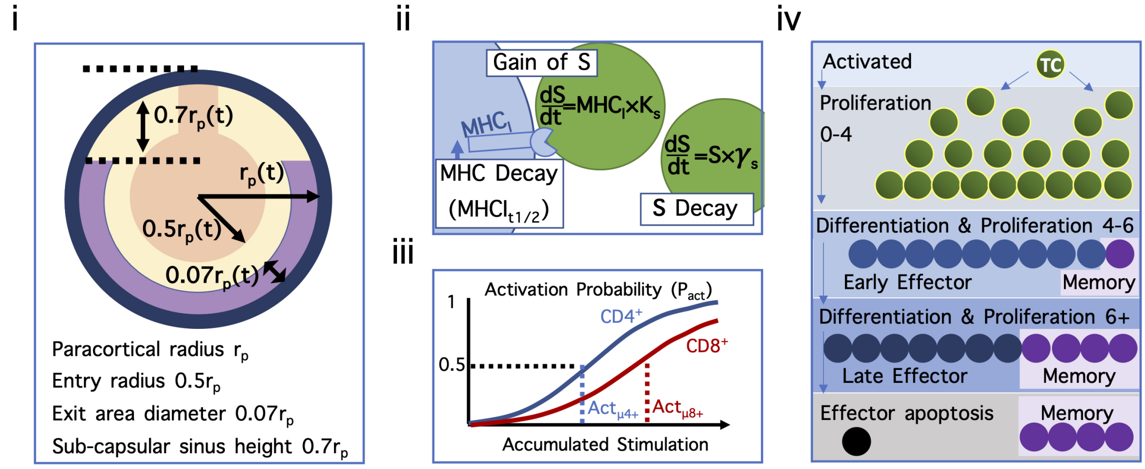

**Fig B Modelling Sphingosine-1-phospate-1 receptor ($S1P_1r$)-mediated retention.** Regulation of $S1P_1r$ on (i) naïve TCs post-transmigration from the blood, (ii) naïve TCs during LN shutdown and (iii) activated and early effector TCs.

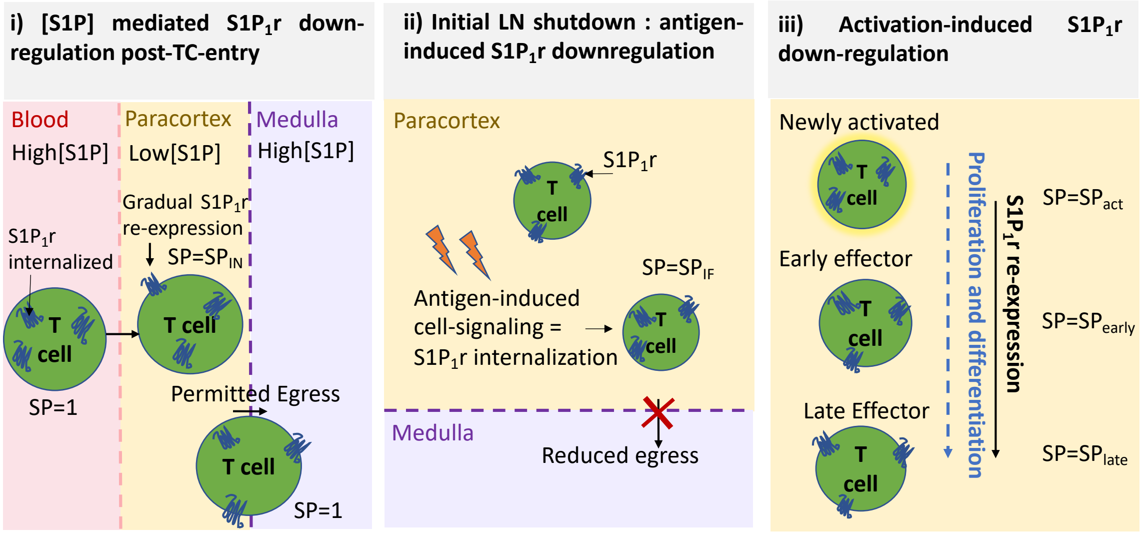

**Fig C A class diagram displaying the underlying ABM structure.** The model is constructed using instructions in the 'context builder' class. The entire modelling domain is described by the context class, and each compartment of the domain is described by the GridCell class. In a 3D simulation, each grid cell can be queried to identify the 26 neighbouring grids and how many agents they contain. The TC class is a template for the TC object produced and is instantiated thousands of times to create TCs with the same variables but slightly different values. A subclass of cognate TCs extends the template with more methods and variables relating to interaction and proliferative response.

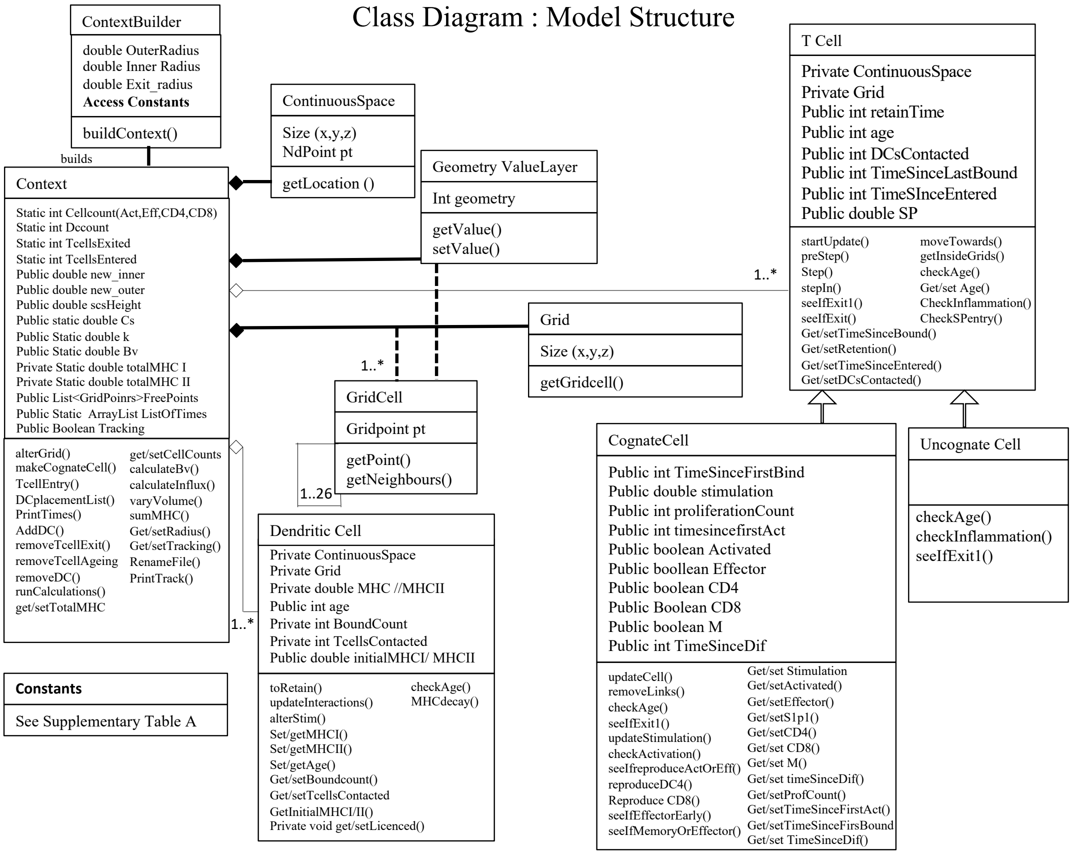

### S1 File Supplementary Parameter Tables

#### Table B Parameters that were not varied in the global sensitivity analysis

| Symbol | Parameter | Value | Reference |
|---|---|---|---|
| | Model Geometry | | |
| $r_p$ | Initial paracortex radius | 200$\mu$m | [44,45] |
| - | Entry radius | 0.5 $r_p$ | [44,45] |
| - | Exit radius | 0.07$r_p$ | [44,45] |
| - | Sub Capsular Sinus height | 0.7$r_p$ | [44,45] |
| GS | Grid Size | 6$\mu$m | - |
| | TC Properties | | |
| - | Initial occupation | 55% | [46] |
| - | Radius | 3.3$\mu$m | [47] |
| - | Ratio CD4:CD8 | 0.7:0.3 | [44,45] |
| - | Lifespan naive | 0.5 $r_p$ | [48] |
| - | Lifespan naive | 0.5 $r_p$ | [49] |
| - | TC entry Afferent:HEV ratio | 0.1:0.9 | [50,51] |
| $\mathrm{Act}l_{4+}$ | Slope of $CD4^+$ activation curve | -69.81 | - |
| $\mathrm{Act}l_{8+}$ | Slope of $CD4^+$ activation curve | -80.71 | - |
| $\mathrm{Dif}l_{4+}$ | Slope of $CD4^+$ differentiation curve | -17.26 | - |
| $\mathrm{Dif}l_{8+}$ | Slope of $CD8^+$ differentiation curve | -13.58 | - |
| | T cell movement | | |
| $\beta$ | Probability of movement | 0.6 | [33,52–55] |
| $P_e$ | Probability of egress | 0.0126 | - |
| $\gamma$ | Max cells per grid | 2 | - |
| $T_{res}$ | TC residence time | 24hrs | [12,56] |
| | DC properties | | |
| - | DC span | 2 grids | [47,57,58] |
| - | DC Lifespan | 2.5days | [1,59,60] |

**Table C Parameters varied in the global sensitivity analysis. Continued overleaf.**

| Symbol | Parameter Description | Default | Min | Max | Mean | SD | Distrib. | Ref |
|---|---|---|---|---|---|---|---|---|
| | TC response parameters | | | | | | | |
| Act$\mu_4$ | $CD4^+$ activation curve mean | 120 | 70 | 230 | - | - | Unif | [61–66] |
| Act$\mu_8$ | $CD8^+$ activation curve mean | 140 | 90 | 250 | - | - | Unif | [61–66] |
| Dif$\mu_4$ | $CD4^+$ differentiation curve mean | 60 | 30 | 90 | - | - | Unif | [61–66] |
| Dif$\mu_8$ | $CD8^+$ differentiation curve mean | 40 | 20 | 60 | - | - | Unif | [61–66] |
| $TP_4$ | Min time between $CD4^+$ proliferations (hrs) | 11 | - | - | 11 | 1.16 | Norm | [29–32] |
| $TP_8$ | Min time between $CD8^+$ proliferations (hrs) | 7 | - | - | 7 | 0.88 | Norm | [29,30] |
| $Max_{P8}$ | Max proliferations $CD8^+$ | 16 | - | - | 16 | 1.2 | Norm | [38,67–69] |
| $Max_{P4}$ | Max proliferations $CD4^+$ | 10 | - | - | 10 | 1.2 | Norm | [30–32] |
| $Dif_{early}$ | Early Memory:Effector cell differentiation | 0.01 | 0.001 | 0.02 | 0.01 | - | Exp | [70] |
| $Dif_{late}$ | Late Memory:Effector cell differentiation | 0.04 | 0.01 | 0.08 | - | - | Unif | [70] |
| | TC interaction dynamics | | | | | | | |
| $T_{NC}$ | Mean non-cognate T-DC interaction (min) | 3.5 | - | - | 3.5 | 1 | Norm | [60,71] |
| $T_{short}$ | Short cognate TC-DC interaction (min) | 10-15 | - | - | 10 | 3 | Norm | [24,60,71] |
| $T_{long}$ | Long cognate TC-DC interaction (min) | 50-70 | - | - | 50 | 12 | Norm | [23–25,72] |
| $T_{change}$ | Time TCs switch to long interactions (hr) | 8 | - | - | 8 | 1 | Norm | [23–25,72] |
| $B_{max}$ | Max TCs a DC can bind per-step | 3 | 1 | 5 | - | - | Unif | - |
| $B_{step}$ | Max TCs a DC can bind | 15 | 4 | 20 | - | - | Unif | [73] |
| | TC Stimulation | | | | | | | |
| $K_s$ | Stim. gain coefficient | 0.015 | 0.005 | 0.02 | - | - | Unif | - |
| $\lambda$ | TC stim. decay factor | 0.99 | 0.99545 | 0.9999 | - | - | Unif | - |
| $MHC_i$ | Initial MHCI/II | 250 | 150 | 350 | - | - | Unif | [18–21] |
| $MHCI_{1/2}$ | MHCI half life (hrs) | 19.7 | - | - | 19.7 | 6 | Norm | [18,19] |
| $MHCII_{1/2}$ | MHCII half life (hrs) | 60 | - | - | 60 | 6 | Norm | [20,21] |
| $F_{cog}$ | Frequency of cognate TCs that enter | 1e-4 | 5e-5 | 1.5e-4 | - | - | Unif | [31,74–76] |
| $\Phi_{DC}$ | Total DCs entering as % of initial TCs | 0.04 | 0.02 | 0.06 | - | - | Unif | [1] |
| $T_{DCin}$ | DC entry duration (days) | 2.5 | 0.5 | 4.5 | - | - | Unif | [1] |

| Symbol | Parameter Description | Default | Min | Max | Mean | SD | Distrib. | Ref |
|---|---|---|---|---|---|---|---|---|
| | Sphingosine-1-phosphate receptor regulation | | | | | | | |
| $SP_{entry}$ | $S1P_1$r expression post entry | 0.1 | 0.01 | 1 | - | - | Unif | [41,42,77] |
| $SP_{act}$ | $S1P_1$r expression when activated | 0.01 | 0.001 | 0.02 | - | - | Unif | [41–43,77] |
| $S1_{early}$ | Effector $S1P_1$r (Proliferation<=6) | 0.4 | 0.01 | 1 | - | - | Unif | [41,43] |
| $SP_{late}$ | Effector $S1P_1$r (Proliferation>6) | 0.8 | 0.3 | 1.3 | - | - | Unif | [41,43] |
| $SP_{mem}$ | Memory $S1P_1 r$r | 1 | - | - | 1 | 0.1 | Norm | [41,43] |
| $SP_{IF}$ | $S1P_1 r$ on all TCs during inflam. | 0.4 | 0.2 | 0.8 | - | - | Unif | - |
| $T_{Entry}$ | Time $S1P_1 r$r is low post-entry (min) | 60 | 13 | 120 | - | - | Unif | - |
| $T_{Inflam}$ | Time to alter $S1P_1 r$ during inflam.(hr) | 4 | 1 | 7.5 | - | - | Unif | - |
| | T cell recruitment | | | | | | | |
| RT1 | recruitment increase stim. threshold | 2e4 | 2e4 | 1e5 | - | - | Unif | [4,6,78,79] |
| RT2 | Stim. threshold for max. recruitment | 4e5 | 2e5 | 2e6 | - | - | Unif | [4,6,78,79] |
| $R_F$ | Recruitment Factor | 3e-6 | 1e-6 | 4e-6 | - | - | Unif | [4,6,78,79] |
| | Paracortex expansion | | | | | | | |
| $V_{Max}$ | Max fold-volume increase | 1.00 | 2.00 | 2.50 | - | - | Unif | - |
| l | Rate of volume change around m | 7e-05 | 3e-05 | 1e-04 | - | - | Unif | - |
| $T_{mid}$ | No. of TCs for 50% max-volume | 120000 | 90000 | 150000 | - | - | Unif | - |
