## Supplementary Files 1-5 for "Lymph node swelling combined with temporary effector T cell retention aids T cell response in a model of adaptive immunity": S2_File.pdf

### Supplementary File 2: Supplementary Methods B

The following section presents the logic underlying the sub-methods called during the simulation. The order that each method is called is depicted in the 'Main' diagram (Fig 2B).

#### 2.1 UML diagrams describing the rules used to build the ABM

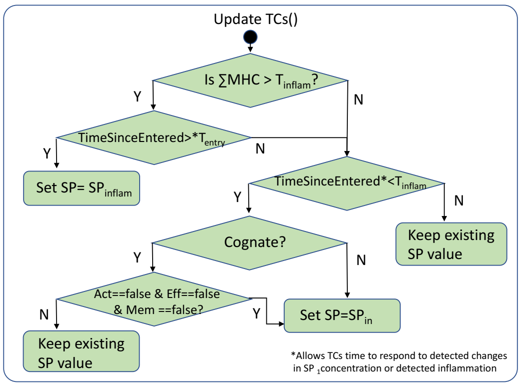


Figure 1: UML diagram for 'UpdateTCs'. By checking the time since the TC entered, a 'lag time' representing time for detection of signals and receptor internalisation to occur.

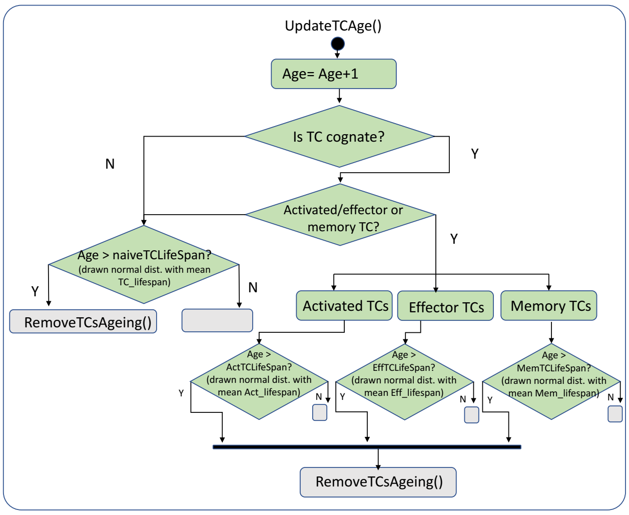


Figure 2: UML diagram for 'UpdateTCsAgeing'.

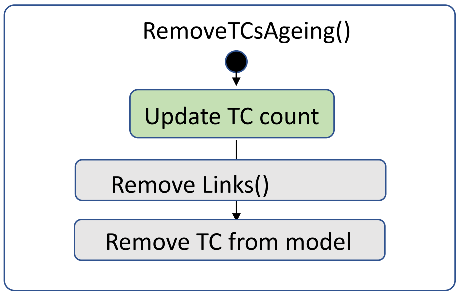


Figure 3: UML diagram for 'RemoveTCsAgeing'. The different types of TCs draw ages from gaussian distributions with a different mean value.

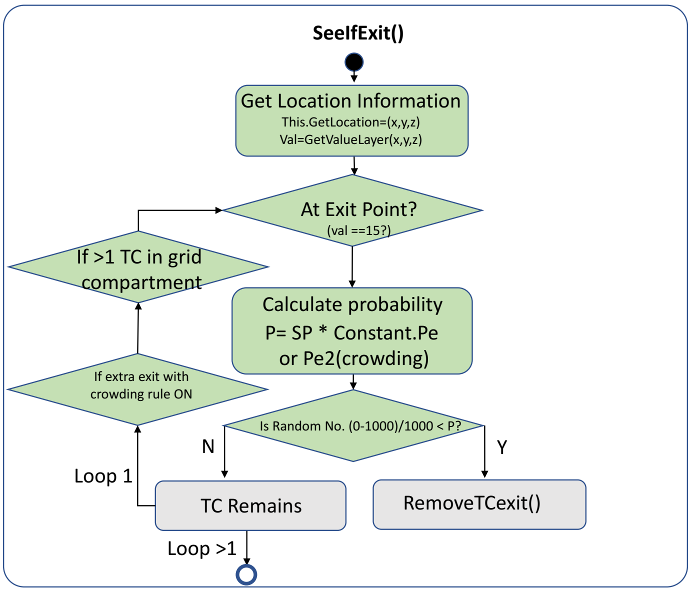


Figure 4: UML diagram for 'SeeIfExit'. The original simulations did not incorporate the second check of TC crowding (crowding rule).

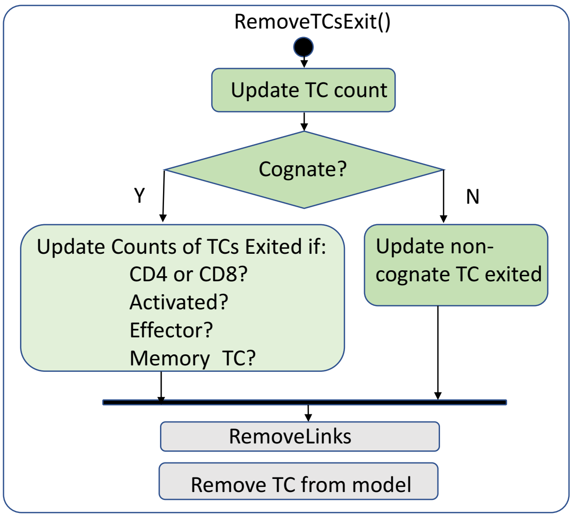


Figure 5: UML diagram for 'RemoveTCExit'. Ageing and TC exit are treated differently due to different counters being updated

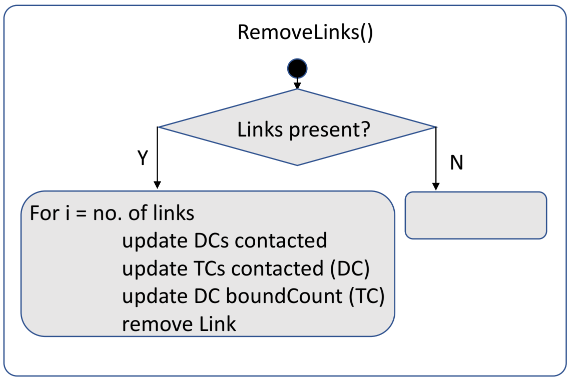


Figure 6: UML diagram for 'RemoveLinks'. Links between TCs and DCs are formed using a projection network, a feature of RepastSimphony, and must be removed when the connection is over.

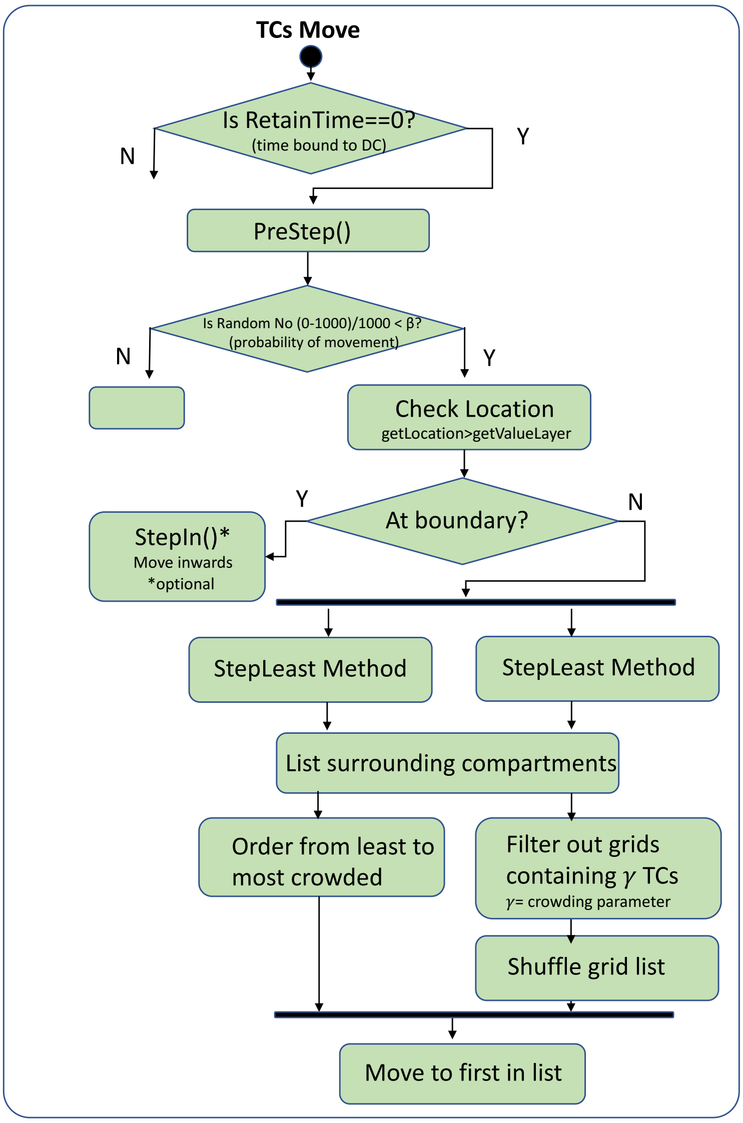


Figure 7: UML diagram for 'TCsMove'. The model has the capacity for different movement options, and maximum TCs per grid is set with crowding parameter $\gamma$

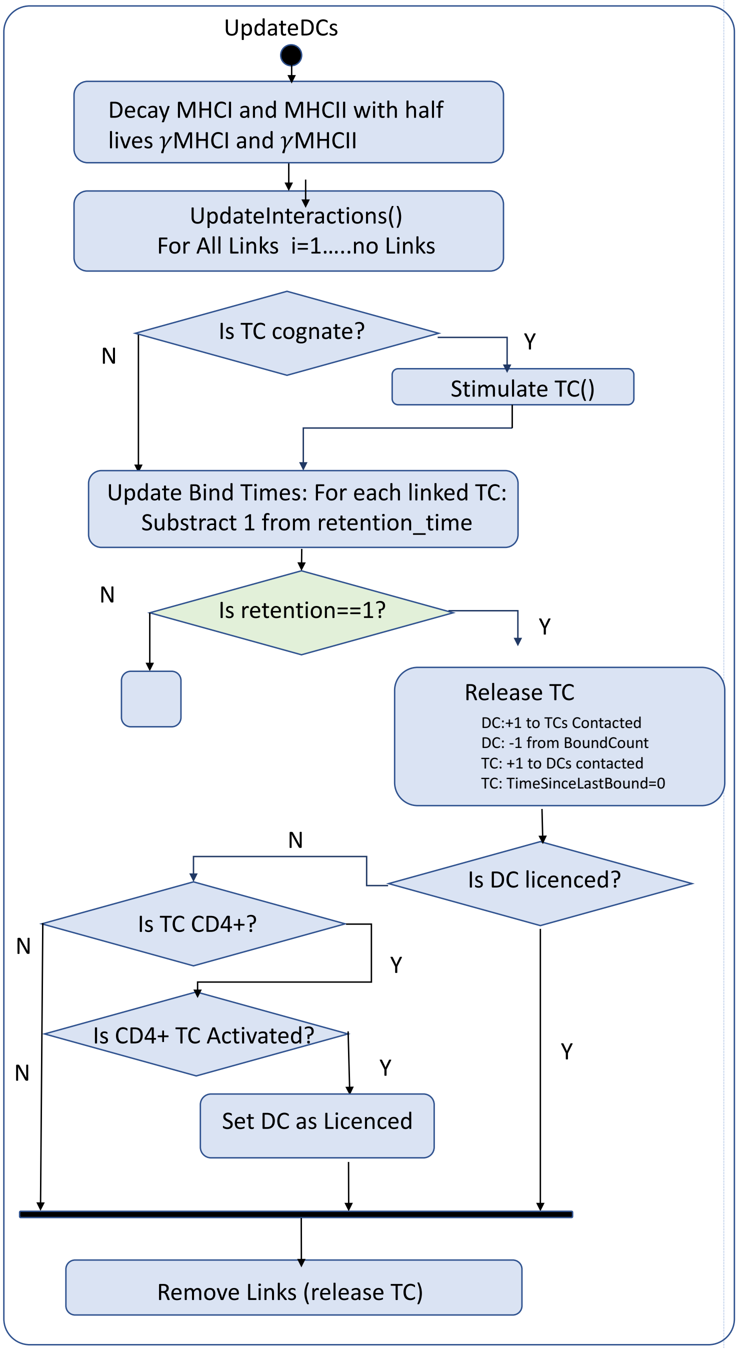


Figure 8: UML diagram for 'UpdateDCs'. DCs enter as mature migrating DCs but may become licenced through interaction with activated CD4+ TCs

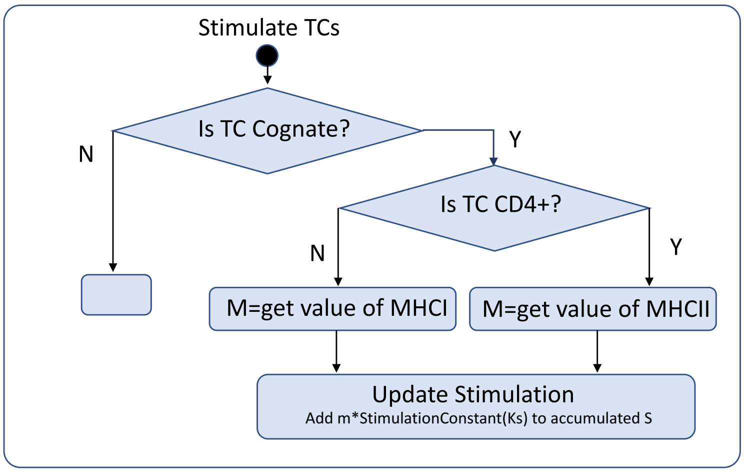


Figure 9: UML diagram for ’StimulateTCs’. DCs can access the variables of the agent attatched via a projection link and thus alter their state based on the DCs MHCI/II presentation and the Stimulation constant, Ks.

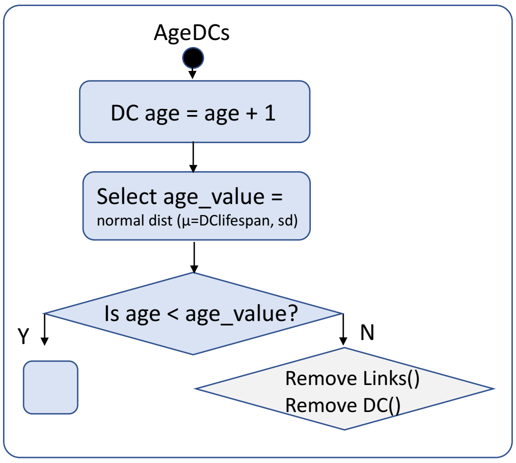


Figure 10: UML diagram for ’AgeDCs’. Mature DCs have a lifespan of around 2.5 days.

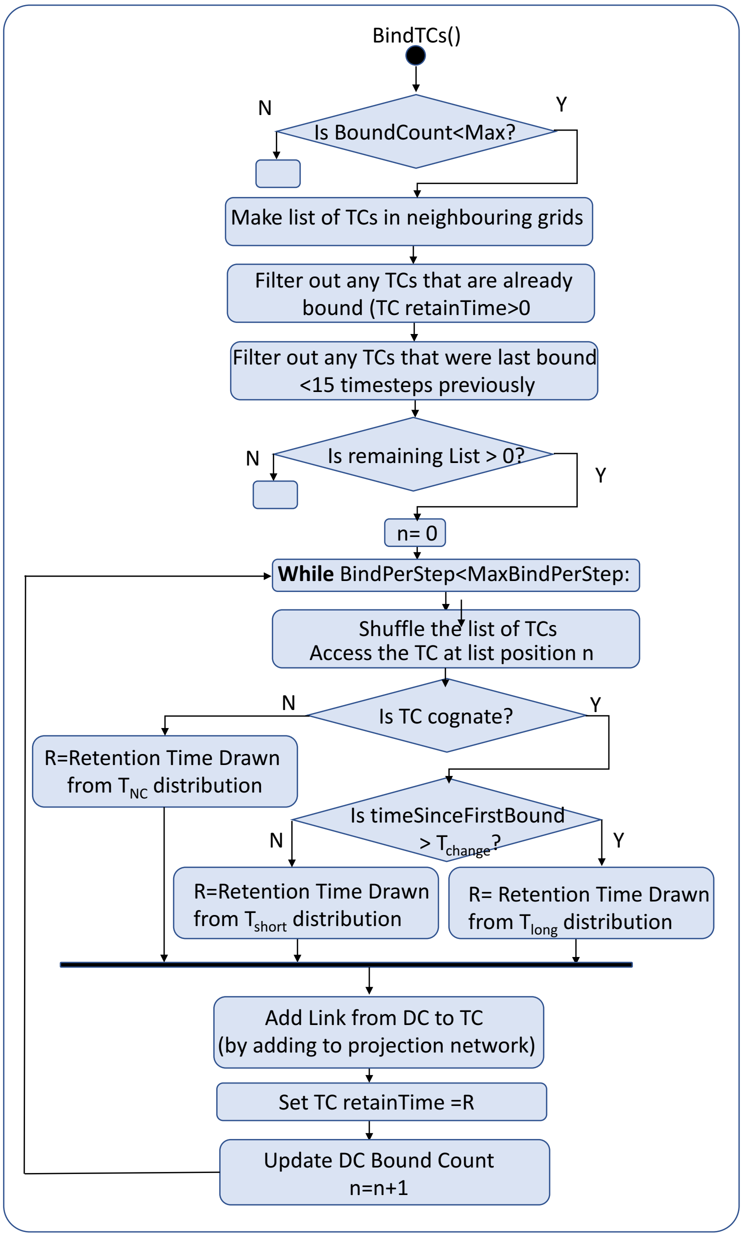


Figure 11: UML diagram for 'BindTCs'. TCs receive a 'interaction time' drawn from a relevant probability density function, based on their type and history.

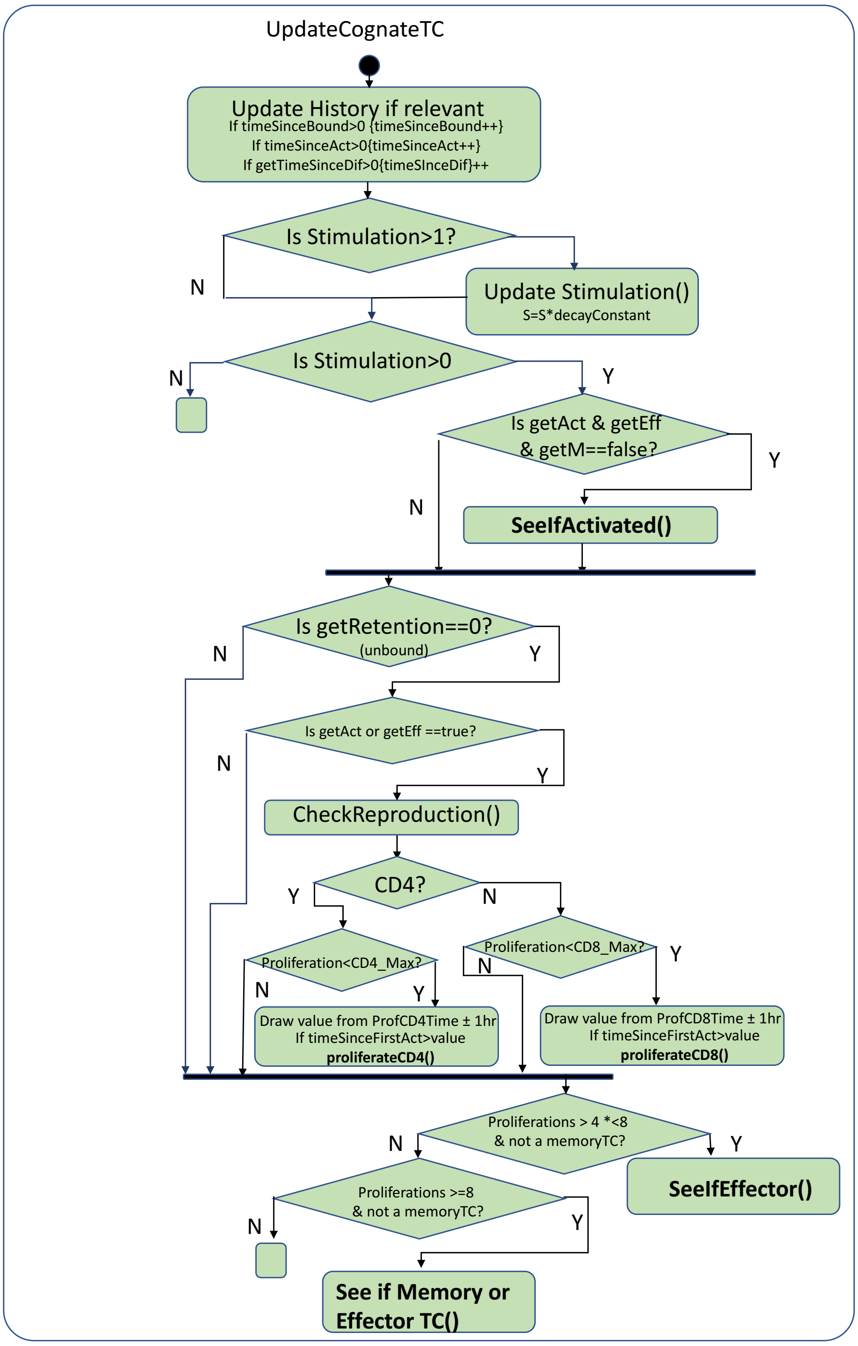


Figure 12: UML diagram for 'UpdateCognateTCs'.

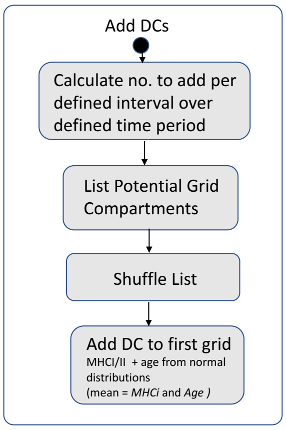


Figure 13: UML diagram for 'AddDCs'.

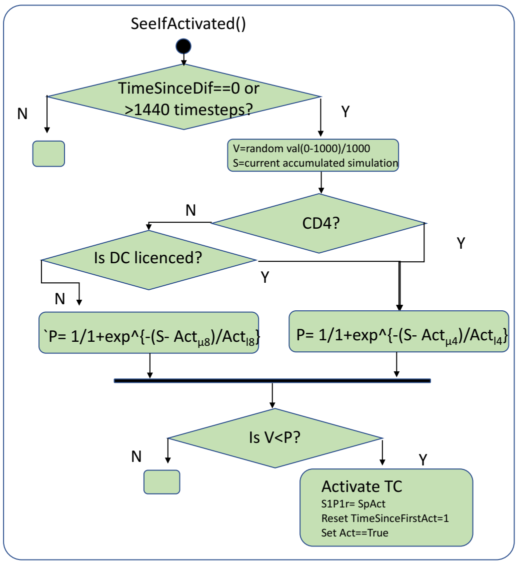


Figure 14: UML diagram for 'SeeIfActivated'.

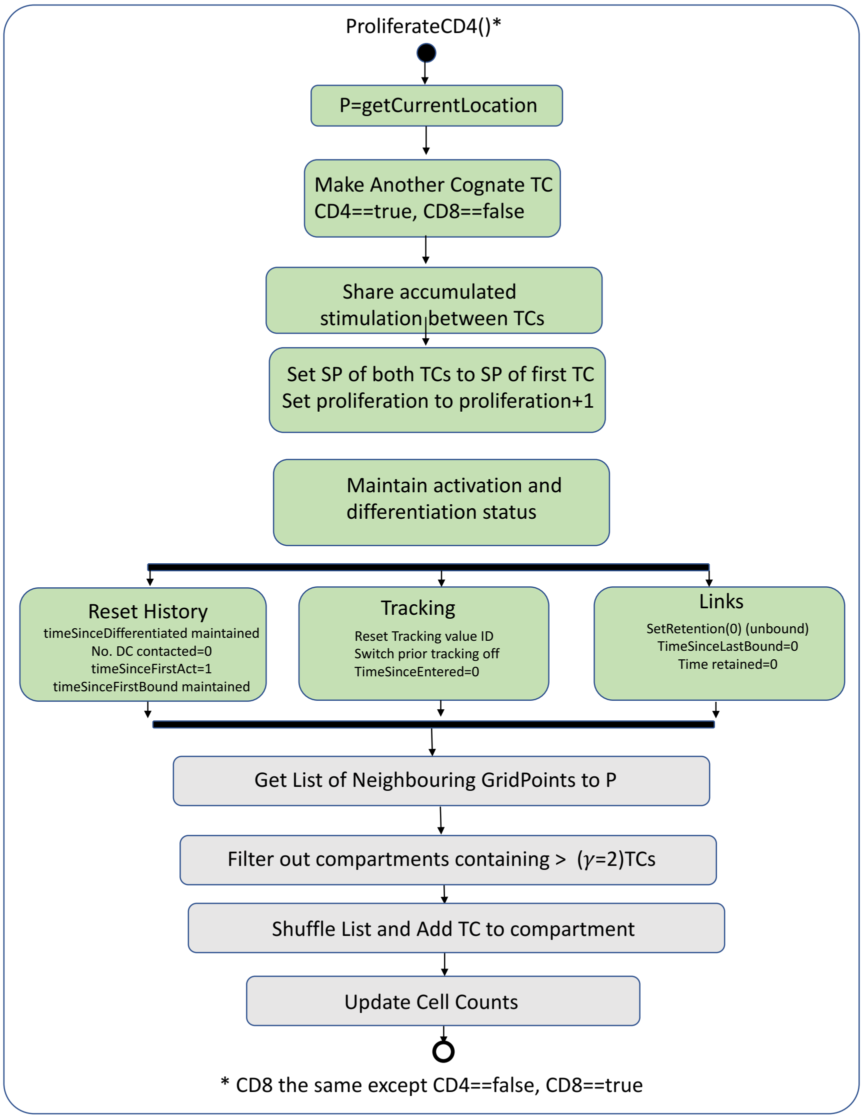


Figure 15: UML diagram for 'Proliferation'.

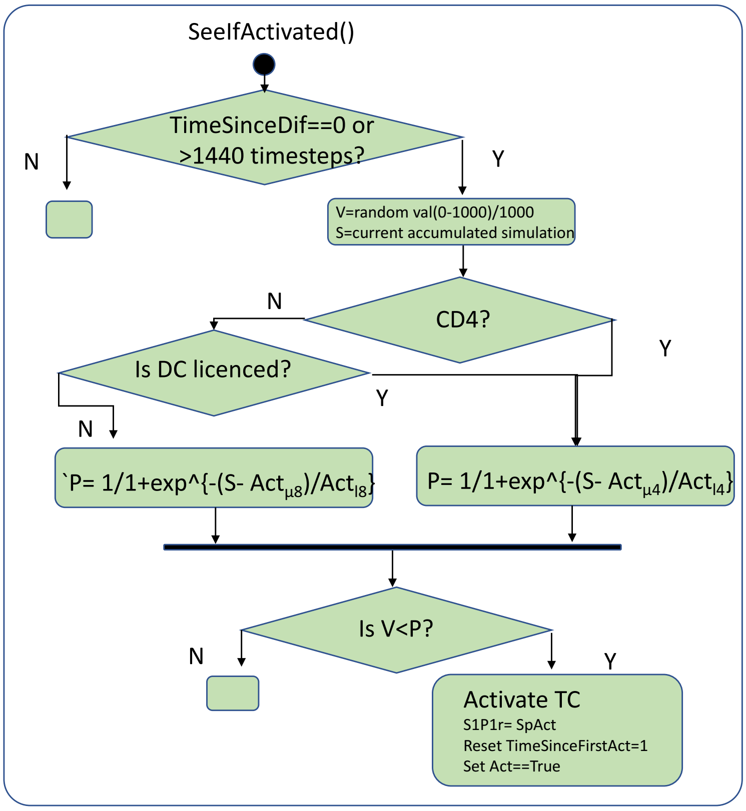


Figure 16: UML diagram for 'SeeIfEffector' or 'SeeIfEffectorMemory. The two behaviours are similar but for seeIfEffectorMemory, the time since differentation cannot be zero, as the TC must have previously differentiated, and more memory TCs are made.

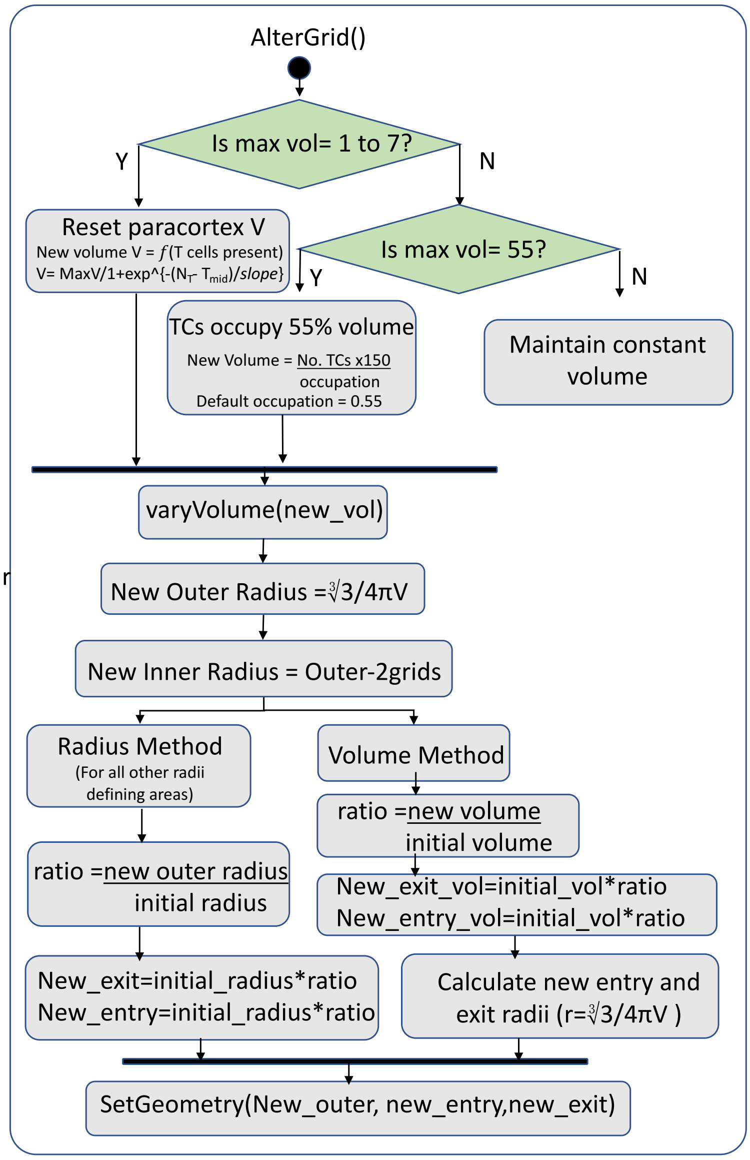


Figure 17: UML diagram for 'AlterGrid'. This methods carries out the calculation of the new paracortex volume then passes the calculated radii to 'Set Geometry'.

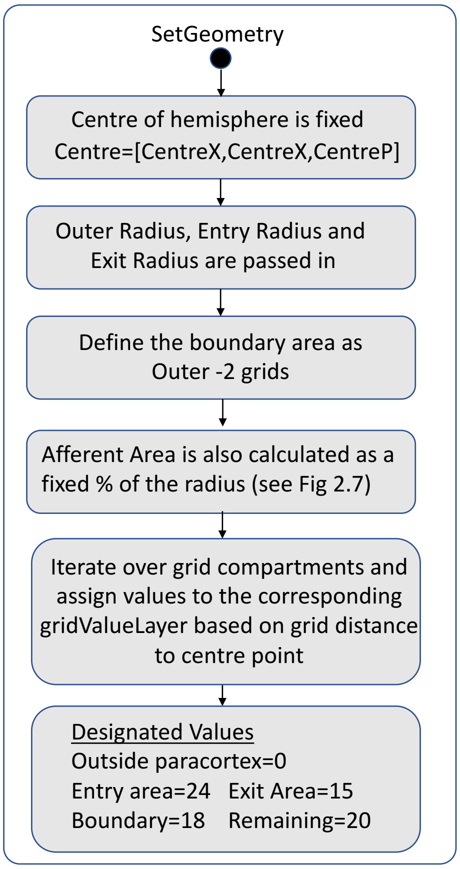


Figure 18: UML diagram for 'SetGeometry'. Only the radii are required as inputs to determine the geometry of the LN
