## Supplementary Files 1-5 for "Lymph node swelling combined with temporary effector T cell retention aids T cell response in a model of adaptive immunity": S3_File.pdf

### Supplementary File 3 : Supplementary Results A

#### Supplementary figures for results from baseline calibration, model validation and LN swelling simulations.

**Fig A Captured phases of TC trafficking and response to AgDC stimuli.** Changes in proliferation and differentiation continued after the initial stimulus was no longer present. TC recruitment and TC egress changes also accompanied the response.

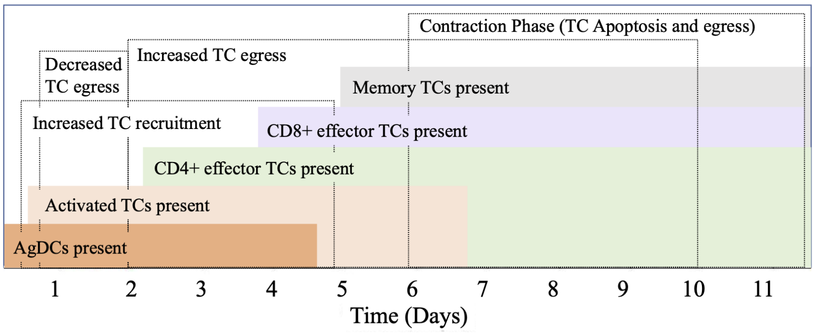

**Fig B TC responses *in-silico* and *in-vivo* when proportion of cognate TCs was varied**. i. The $CD4^+$ magnitude of response in the dLNs of mice post-antigen injection correlated to estimated starting frequency of cognate TCs in a sample of $1x10^7$ TCs [1]. ii. Results *in-silico* showed an overall increase in cognate $CD4^+$ TC response with increasing $F_{cog}$ (n=8). Simulations using a wider range of $F_{cog}$ values (iii, iv) confirmed total cognate $CD4^+$ TCs and $CD8^+$ TCs increased linearly with $F_{cog}$. v. Mice were infected with VSV-M45 or VSV-ova with starting precursor $CD8^+$ frequencies of $7x10^{-5}$, $8x10^{-5}$ and $13x10^{-5}$ respectively. The peak number of resulting TCs as a percent of overall $CD8^+$ TCs present is shown [2]. vi. Simulations using the same $F_{cog}$ *in-silico* showed a similar increasing trend in cognate $CD8^+$ TCs with similar increase rate.

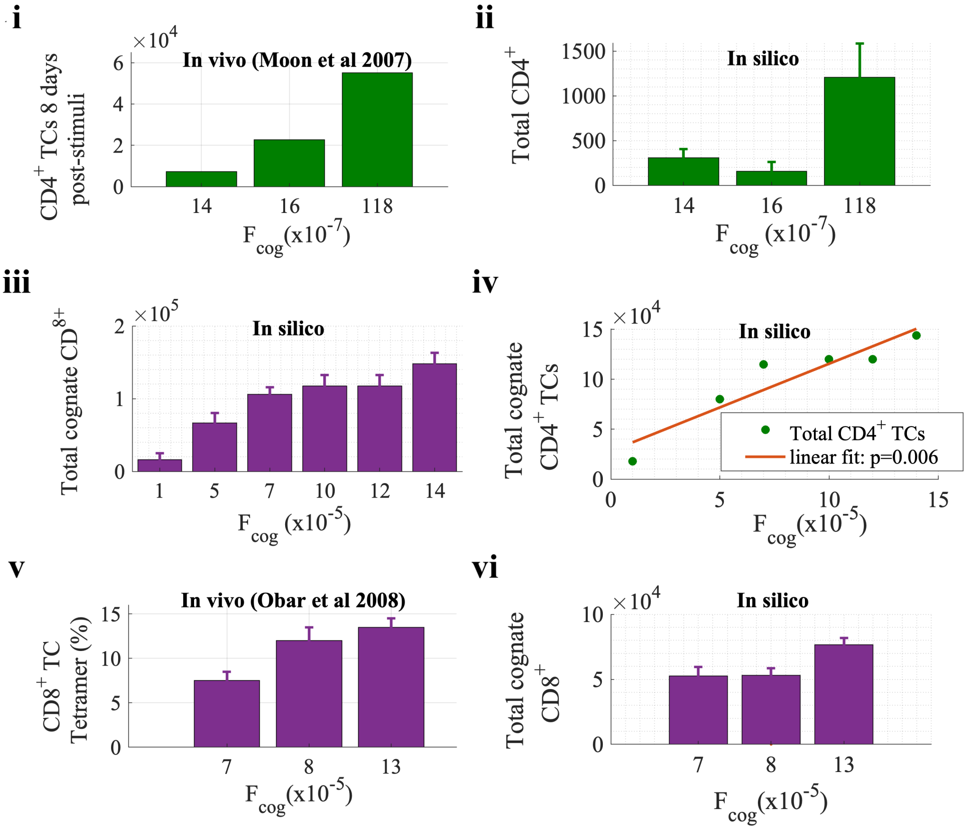

**Fig C TC responses *in-silico* and *in-vivo* when a stimulus is abruptly abolished.** i. The LNs of transgenic rats were injected with OT-1 CD8$^+$ TCs specific for agDCs that were subsequently injected. The lifespan of the agDCs were curtailed using DT injection at 1hr and 12hrs and the change in subsequent CD8$^+$ effector TC response recorded. Adapted from Prlic et al 2006 [3]. ii. The simulated disrupted input stimuli achieved by curtailing the 60hr DC influx at 12±2hr. iii-iv. The results of simulations (n=8). Mean (±SEM) CD8+ TCs were reduced 91% when the stimulus was curtailed at 12hrs compared to sustained entry for 60hrs. iv. individual CD8$^+$ TC responses varied by a factor of 10 in an all or nothing response manner.

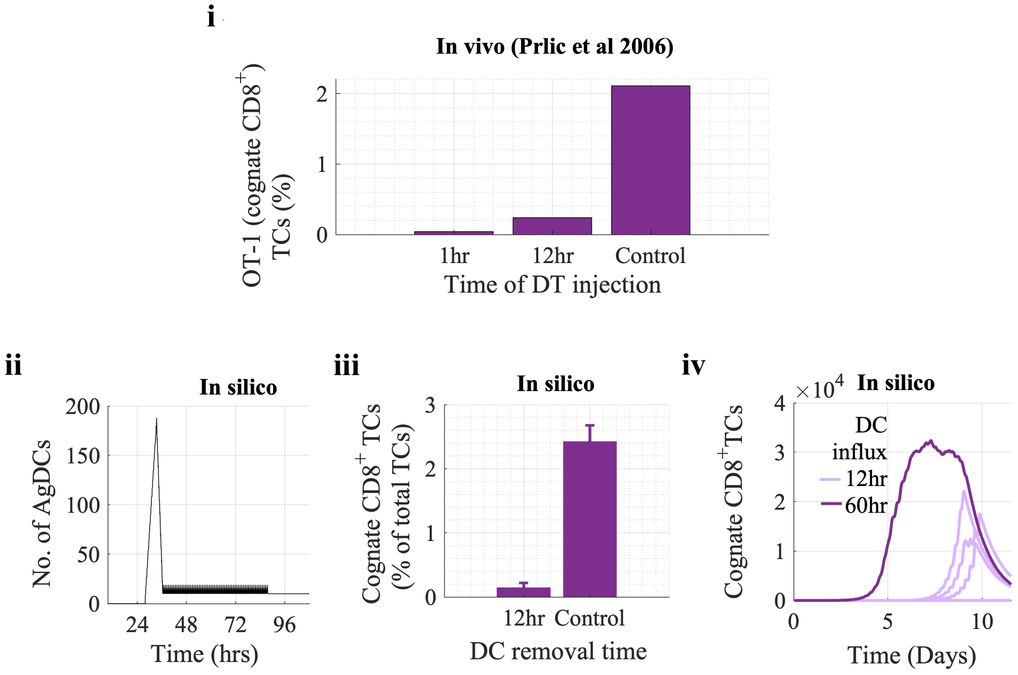

**Fig D TC responses *in-silico* and *in-vivo* when the DC stimuli (no. of DCs applied as a fraction of initial TCs present, ($\phi$DC)) is varied.** The estimated percentage of CD8$^+$ TCs that underwent a proliferative response (i) in the LNs of chimeric mice 7 days post-injection with antigen LM-GP33 and (ii) in cell culture post-application of DCs [4]. (iii) Analysis of *in-silico* CD8$^+$ TC response at low doses showed a significant response (1x10$^4$ total cognate CD8$^+$ TCs) or no proliferative response at all (<10 cognate CD8$^+$ TCs). (iv) *In-silico* simulations increasing the proportion of DCs resulted in increasing numbers of cognate CD8$^+$ TCs that plateaued as the DC dose increased.

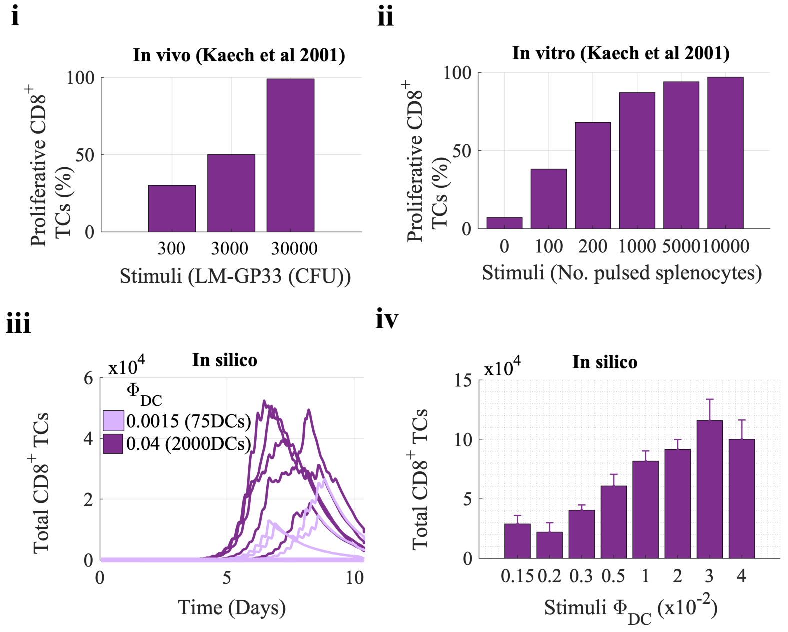

**Fig E Supplementary results when varying maximal swelling ($V_{max}$) and TC number required to reach half $V_{max}$ ($T_{mid}$).** With default $T_{mid}$=$10^5$ TCs, no correlation was observed between $V_{max}$ and (i) effector TCs exited, (ii) total cognate $CD4^+$ TCs and (iii) total cognate $CD8^+$ exited. (iv) When $T_{mid}$ was increased to 1.2 x$10^5$ TCs, TC activation increased with maximal swelling ($R^2$=0.96, p=9.67x$10^{-6}$), (v) cognate $CD4^+$ TCs exited weakly correlated with $V_{max}$ ($R^2$=0.71,p=0.002) and (vi) cognate $CD8^+$ TCs exited showed no significant correlation. (vii) Increasing ($V_{max}$) from 1.2 to 2.0 decreased effector TC number by 17% and to 2.5 by 5%. (viii) Activation of TCs increased with $V_{max}$ but negatively correlated with $T_{mid}$ ($R^2$=0.78, 0.99, 0.84, 0.84 for $V_{max}$=1.2, 1.5 , 2.0 and 2.5 respectively, p=1.5x$10^{-5}$, 1.5x$10^{-5}$, 0.01, 0.01).

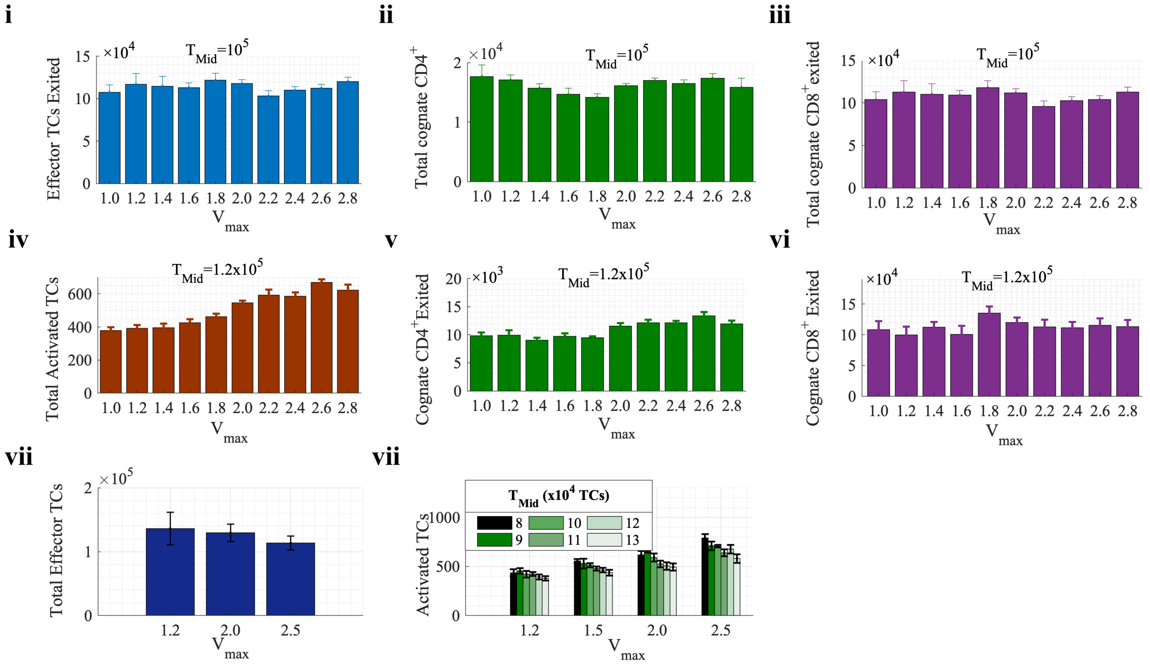
