## Supplementary Files 1-5 for "Lymph node swelling combined with temporary effector T cell retention aids T cell response in a model of adaptive immunity": S4_File.pdf

### Supplementary File 4: Supplementary Results B

#### 4.1 Global Sensitivity Analysis Results

During the global sensitivity analysis, the partial rank correlation coefficient between parameters and output of interest were calculated at each day of the simulation. The results indicate that TC activation was most sensitive to the frequency of TC recognition ($F_{cog}$), stimuli strength ($\phi_{DC}$) and stimuli timing ($T_{DCin}$) throughout week 1. Activation of TCs was also sensitive to the recruitment factor ($R_F$) at day 2. Sensitivity to maximal swelling ($V_{max}$) was also identified between days 2 and 4, showing a positive correlation with TC activation (Fig A and Table A-C in S3 File).

Parameters that would drive TC response, such as $F_{cog}$ and ($\phi_{DC}$), displayed a positive correlation with effector TC present and effector TCs exited in the first week only. Maximal swelling became a highly influential factor in week 2, displaying a strong negative correlation with effector TC production. During week 2, the recruitment factor (and therefore TC recruitment) and the maximum number of CD8+ proliferations also showed a negative correlation. It is possible that the positive influence of $F_{cog}$ and ($\phi_{DC}$) is lost, and TC recruitment and proliferation becomes negative as increased TC number also drives swelling and TC egress too early. A key reason for carrying out the global sensitivity analysis was to ensure that potentially sensitive uncertain parameters, which could significantly influence outcomes of interest, were identified. However, the unconstrained parameters used to describe signal integration, such as mean amount of signal accumulated to activate or differentiate, were not identified as sensitive relative to the other parameters.

**Fig A The sensitivity of output (i) the activated TCs present or (ii) the effector TCs present, to parameters over the time course of the simulation.** The Partial Rank Correlation Coefficient (PRCC) between parameter and output measure is displayed, with the grey area representing weak PRCC values that are not significantly different from zero.

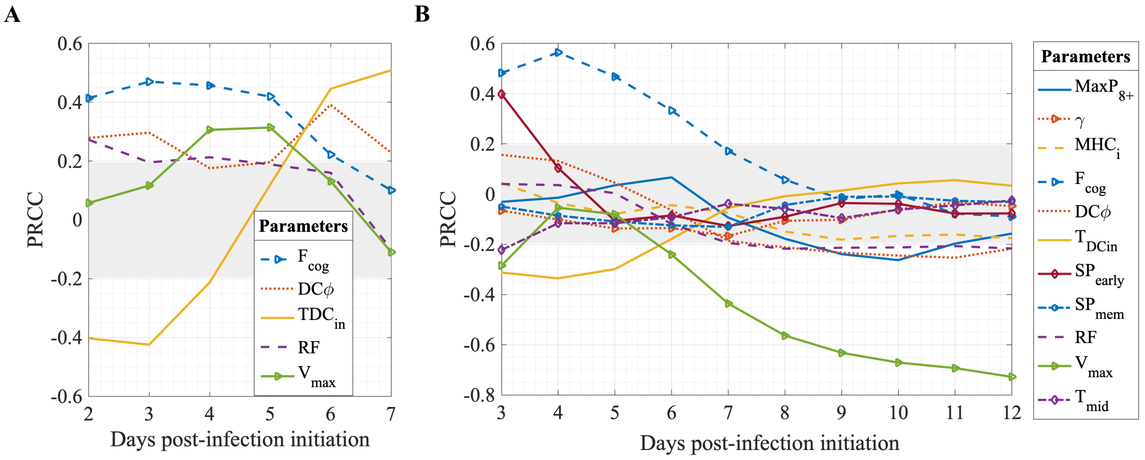

#### S4 File Supplementary Tables

**Table A Parameters that significantly affected the number of activated TCs in the paracortex from day 2 to 6 post-initiation of infection.**

| Parameter | Day2 | Day 3 | Day 4 | Day 5 | Day 6 |
|---|---|---|---|---|---|
| $F_{cog}$ | +++ | +++ | +++ | +++ | ++ |
| $\phi_{DC}$ | +++ | ++ | +++ | | |
| $T_{DCin}$ | −−− | −−− | ++ | +++ | +++ |
| $R_F$ | | +++ | +++ | | − |
| $V_{max}$ | + | +++ | +++ | | − |

Key : +/-= 0.05> $p$ >0.001  ++/− =0.001> $p$ >$10^{-6}$  +++/— = p< $10e^{-6}$

**Table B Parameters that significantly affected the (top) effector TCs present in the paracortex and (bottom) effector TCs that exited the paracortex from day 3-12 post-initiation of infection.**

| Parameter | Day 3 | Day 4 | Day 5 | Day 6 | Day 7 | Day 8 | Day 9 | Day 10 | Day 11 | Day 12 |
|---|---|---|---|---|---|---|---|---|---|---|
| $Max_{P8+}$ | | | | | − | −− | −− | −−− | −− | − |
| $Max_{P4+}$ | | | | + | | | | | | |
| $T_{NC}$ | | | | | | | | | + | + |
| $T_{short}$ | + | | | | | | | | | |
| $MHC_i$ | | | | | | | | | | − |
| $F_{cog}$ | +++ | +++ | +++ | +++ | | | | | | |
| $\phi_{DC}$ | +++ | +++ | + | | | | | | − | − |
| $T_{DCin}$ | −−− | −−− | −−− | −− | | | | | | |
| $SP_{early}$ | | − | | | | | | | | |
| $R_F$ | | | | | | − | −− | −− | −− | −− |
| $V_{max}$ | | ++ | | + | −−− | −−− | −−− | −−− | −−− | −−− |

| Parameter | Day 3 | Day 4 | Day 5 | Day 6 | Day 7 | Day 8 | Day 9 | Day 10 | Day 11 | Day 12 |
|---|---|---|---|---|---|---|---|---|---|---|
| $Max_{P8+}$ | | | | | | −− | −− | −− | − | |
| $\gamma$ | | | − | − | − | | | | | |
| $MHC_i$ | | | | | | | −− | −− | − | − |
| $F_{cog}$ | +++ | +++ | +++ | +++ | + | | | | | |
| $\phi_{DC}$ | + | | | | −− | −− | −− | −− | −− | −− |
| $T_{DCin}$ | −−− | −−− | −− | − | | | | | | |
| $SP_{early}$ | +++ | | | | | | | | | |
| $SP_{mem}$ | | | | | − | | | | | |
| $R_F$ | | | | | −− | −− | −− | −− | −− | −− |
| $V_{max}$ | −− | | | −− | −−− | −−− | −−− | −−− | −−− | −−− |
| $T_{mid}$ | −− | | | | | | | | | |

Key : +/-= 0.05> $p$ >0.001  ++/− =0.001> $p$ >$10^{-6}$  +++/— = p< $10e^{-6}$

**Table C Parameters that significantly affected the number of (left) memory T cells present and (right) number of memory TCs exited.**

| Parameter | Day5 | Day 7 | Day 9 | Day 12 |
|---|---|---|---|---|
| $Max_{P8+}$ | | ++ | +++ | + |
| $Diff_{late}$ | +++ | ++ | | + |
| $\gamma$ | − | | | |
| $F_{cog}$ | +++ | | | − |
| $\phi_{DC}$ | | | | − |
| $T_{DCin}$ | −−− | −− | | |
| $R_F$ | | −− | −− | −− |
| $V_{max}$ | + | − | −−− | −−− |
| $T_{mid}$ | − | | | |

Key : +/-= 0.05> $p$ >0.001

++/– =0.001> $p$ >$10^{-6}$

+++/— = p< $10e^{-6}$

| Parameter | Day5 | Day 7 | Day 9 | Day 12 |
|---|---|---|---|---|
| $TP_{4+}$ | | + | + | |
| $Max_{P8+}$ | | − | −−− | −− |
| $Diff_{early}$ | ++ | | | |
| $Diff_{late}$ | +++ | + | ++ | + |
| $T_{NC}$ | | | | + |
| $\gamma$ | −− | −− | | |
| $MHC_i$ | | | − | − |
| $F_{cog}$ | +++ | | | |
| $\phi_{DC}$ | | −− | −− | −− |
| $T_{DCin}$ | −−− | | | |
| $SP_{early}$ | −−− | − | | |
| $R_F$ | | − | −− | −− |
| $V_{max}$ | | −−− | −−− | −−− |
