## Supplementary Files 1-5 for "Lymph node swelling combined with temporary effector T cell retention aids T cell response in a model of adaptive immunity": S5_File.pdf

### Supplementary File 5: Supplementary Results C

#### Alternative Models

##### 5.1 Consideration of an alternative model with linear TC to LN volume relationship

Implementation of an alternative model of LN swelling, where LN volume was specified as a linear function of TCs present, whilst maintaining TC recruitment rules resulted in a sharp peak in TC number and paracortical volume that was not in-line with in-vivo observations (Fig A). This suggests further rules to represent factors that constrain LN swelling are required, and that driving swelling as a sigmoidal function of TCs present results in expansion that more closely resembles in-vivo observations. Despite the large increase in volume (>7-fold) and TC entry rate, as well as increased TC activation, CD8+ effector TC production is reduced when compared to a moderate doubling of LN volume in a model following the sigmoidal function. The results support our findings that increased swelling aids activation but not effector TC production.

**Fig A TC response when paracortical volume expands as a linear function of the TC count.** Compared to when paracortical volume altered as a sigmoidal function of TC count, with a maximal 1.4-fold or 2.6-fold increase, the (i) maximal paracortical radius more than doubled. (ii) The number of activated TCs doubled. (iii-iv) The total number of effector TCs present and exited was reduced. (v) Peak $CD8^+$ TCs is reduced by at least 35% while (vi) peak $CD4^+$ number is increased by at least 20%.

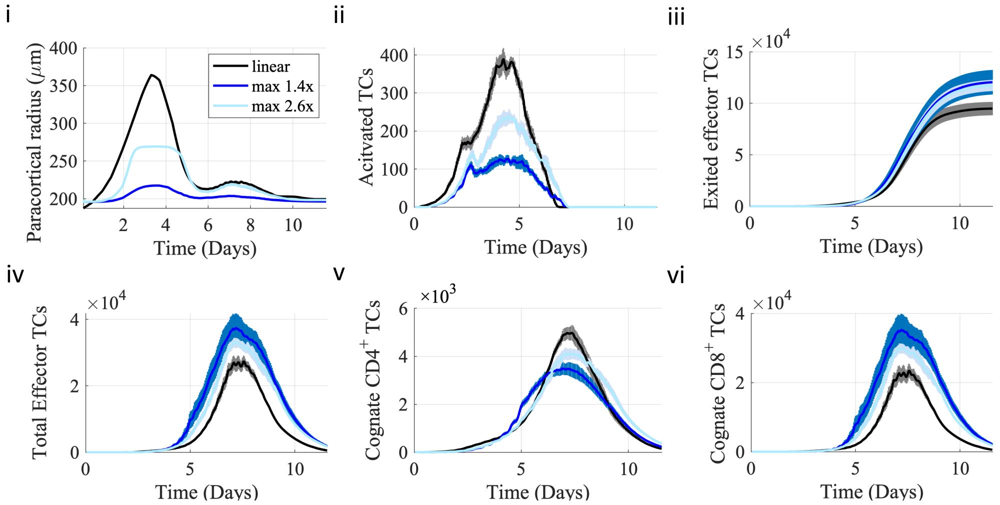

#### 5.2 Considering alternative models of TC crowding and egress

We also noted that with our default modelling assumptions, the average TC to grid compartment ratio fluctuated between 0.55 and 1.3 when swelling was permitted and up to 1.75 when swelling was not permitted. We considered the potential effect of increased TC crowding on exit likelihood, by assuming that when TCs shared an exit grid compartment, it would create additional opportunities to exit, for example due to possible increased pressure inside the node. This only resulted in a significant difference when swelling was not permitted and a $V_{max}$ of 1.8 therefore no clear conclusion was reached (Fig B.ii).

Finally, we implemented a model with constrained growth of exit area with swelling by maintaining the thickness of the cap-shaped area that TCs could exit in. With this constraint, which still allowed some growth in exit area, the number of effector TCs produced increased compared to baseline, but above a 2-fold swelling, the paracortex remained swollen (Fig B.iv). These results suggest that effector TC production in the default swelling model is not limited by lack of DC migration, as effector TC production increase was found at all values of $V_{max}$ , but also that our initial assumption regarding exit areas permitted a return to homeostasis through increased TC egress .

**Fig B The effects of considering alternative models of egress with swelling.** (i) Under baseline modelling conditions, when swelling was permitted, the TC to grid compartment ratio remained between 0.6 and 1.3. Compared to default simulations, effector TC response (ii) was significantly decreased at $V_{max}$ = 1.0 and 1.8 only, resulting in no correlation with Vmax when increased egress with crowding was applied. Modelling lymph node swelling while constraining the expansion of the exit areas by maintaining the thickness of the exit area resulted in (iii) a lack of return to homeostasis if the node swelled more than 1.4x but also meant that (iv) effector TC production increased with sustained swelling at all values of $V_{max}$.

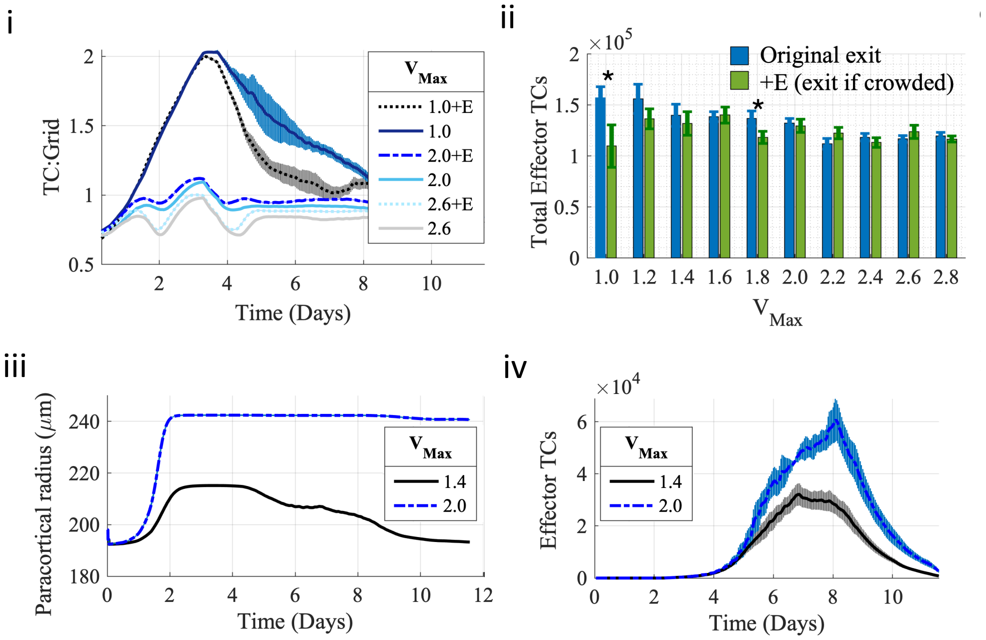
